# HARICOT: De Novo Assembly of Raw Nanopore Signals using Fuzzy de Bruijn Graphs

**DOI:** 10.64898/2026.11.11.765872

**Authors:** Ulysse McConnell, Quang Ngo, Can Firtina

## Abstract

Nanopore sequencing enables reading long nucleotide sequences by measuring characteristic changes in ionic current as molecules pass through a nanopore. These current measurements, known as raw signals, can be analyzed without translating them into bases (i.e., without basecalling), which provides fast and resource-efficient genome analysis. Prior work, Rawsamble, enables finding overlaps between raw nanopore signals, without basecalling them, and generating assemblies from these overlaps with off-the-shelf assemblers such as miniasm. Constructing assemblies directly from raw signals can substantially reduce redundancy and fragmentation in signal datasets, which is useful for many downstream tasks such as constructing non-redundant and accurate basecalled reads from assembled signals. However, existing overlap-based signal assembly strategies suffer from limited scalability and resolution mainly due to using off-the-shelf assemblers (e.g., miniasm) that are designed for basecalled sequences. While scalability and performance limitations can be addressed by existing de Bruijn graph approaches to assembly, their reliance on exact matches makes these methods too sensitive to the inherently noisy raw nanopore signals.

To improve assembly of raw signals using de Bruijn graph approaches, we present HARICOT, a mechanism that constructs assemblies from raw nanopore signals by utilizing fuzzy de Bruijn graphs and Hough-transform-based overlap detection to robustly tolerate errors and noise in raw nanopore signals. Our evaluations on various genomes of different sizes show that HARICOT significantly improves assembly contiguity and quality compared to Rawsamble, helping close the gap to popular assembly tools such as Flye or wtdbg2 operating on nucleotide sequences. At the same time, performance remains comparable to these assemblers, without requiring additional basecalling. We discuss that assembling raw nanopore signals before basecalling enables future work in various ways such as building these de Bruijn graphs in a streaming way and assembly-aware basecalling techniques that can basecall assembly graphs for accurate and redundancy-resolved basecalling.

HARICOT is available at https://github.com/STORMgroup/HARICOT. We also provide the scripts to fully reproduce our results on our GitHub page.

## 1. Introduction

Nanopore sequencing generates readouts of molecules at a high throughput as they pass through a *nanopore* [1]. The presence of a molecule disrupts an ionic current through the pore, which is recorded as a *raw signal* and further processed. Nanopore sequencing offers several advantages as a sequencing technology, including the ability to sequence ultralong reads of up to several Mbp in length [2, 3], perform portable sequencing [4–6], analyze reads in real-time [7–9], and sequence reads selectively [7, 10, 11].

To analyze sequences, raw signals are conventionally first translated into nucleotide sequences by *basecalling* [12–14], which often relies on computationally expensive AI methods such as transformer models [15–17]. It is however possible to analyze sequences without basecalling, an approach called *raw signal analysis* [8, 9, 11, 18–24]. Most existing raw signal analysis methods, such as read mapping [8,9,11,18,19], require a reference genome. Constructing assemblies directly from raw signals [20] removes the need for a reference genome. An assembly of raw signals shows which parts of the raw signals represent each region of the genome, so that only these parts need to be basecalled, as we discuss in Section 4.2.

*De novo* assembly (i.e., assembling a genome from reads without a reference) is an important step in the genome analysis pipeline. Most assemblers can be categorized into one of two paradigms [25, 26]. In *overlap-based* methods, including overlap–layout–consensus (OLC) and string graphs [27–30], nodes correspond to reads and edges to overlaps between them. Approximately matching reads provides more robustness to errors, but requires detecting overlaps between all pairs of reads. In *de Bruijn graph* (dBG) methods [31–34], nodes correspond to *k*-mers, and two nodes are joined by an edge when a (*k* + 1)-mer has one *k*-mer as its prefix and the other as its suffix. Because dBGs rely on exact *k*-mer matches, they can be built more efficiently by counting instead of pairwise matching of reads, but this also makes them sensitive to noisy reads, where a single error can corrupt up to *k* consecutive *k*-mers.

With the emergence of accurate hash-based seeding mechanisms for raw nanopore signals, assemblies from overlapping nanopore signals can be generated [20] using off-the-shelf assemblers such as miniasm [35]. While constructing assemblies from raw signals can be useful to reduce storage requirements by reducing redundancy or designing specialized basecallers for assembled signals, the state-of-the-art raw signal overlapping tool, Rawsamble, produces fragmented assemblies and scales poorly.

These limitations are mainly due to two reasons. First, miniasm is designed for overlaps between basecalled reads and not for overlaps between error-prone raw signals. Second, Rawsamble needs to perform costly chaining between pairs of signals. De Bruijn graph methods can help address scalability issues. However, it is challenging to utilize the existing approaches as 1) de Bruijn graphs require exact *k*-mer matches making them sensitive to noise, 2) noise-tolerant variants such as fuzzy de Bruijn graphs [36] determine approximately matching regions from pairwise overlaps, which are difficult to reliably detect between noisy raw signals, and 3) existing graph cleaning and layout steps are not adapted to the error profile of raw signal assemblies.

Our goal is to assemble raw signals accurately and in a scalable manner. To achieve this, we present HARICOT, a raw signal assembler based on fuzzy de Bruijn graphs. The key idea of HARICOT is to find overlaps while tolerating the noise in raw signals, and to use these overlaps to collapse the parts of different reads that come from the same genomic region into the same node, in three steps. First, to find overlaps *without* connecting seed matches with a costly chaining algorithm [9, 37], HARICOT uses the Hough transform [38], a common technique in computer vision to quickly detect lines among noisy points. Second, to reject false overlaps, HARICOT uses random forest classifiers [39] that take into account various characteristics of the overlap. Third, to remove the remaining errors, HARICOT cleans the assembly graph and joins broken contigs with steps adapted to raw signal assemblies.

Based on our comprehensive analysis using 14 bacterial and eukaryotic datasets, we show that HARICOT 1) is able to generate long contiguous assemblies, with the longest contigs reaching several million basepairs, and 2) performs similar to existing nucleotide assembly tools.

### This paper makes the following key contributions

- We propose HARICOT, the *first* raw signal assembler based on fuzzy de Bruijn graphs, with overlap detection based on the Hough transform and overlap decision based on learned classifiers.
- We show that the longest contigs of HARICOT are on average 15.6× (up to 72×) longer than those of Rawsamble, the only previously published raw signal assembler, and that HARICOT runs on average 3.9× (up to 11.6×) faster than Rawsamble.
- We show that HARICOT produces contigs spanning the entire chromosome on five of six bacterial chromosomes, as Flye [40, 41] does from basecalled reads. On five of seven plant and fly chromosomes, HARICOT constructs more contiguous assemblies than wtdbg2 [36] in terms of NG50. HARICOT provides performance comparable to Flye, even though their elapsed times do not include basecalling.
- We provide the source code, the trained models and the scripts to fully reproduce our results at https://github.com/STORMgroup/HARICOT.

## 2. Methods

HARICOT constructs assembly graphs from noisy raw nanopore signals by tolerating noise via fuzzy de Bruijn graphs.

We provide an overview of the fuzzy de Bruijn graph in Supplementary Section S1. HARICOT operates in five main steps, which are outlined in Figure 1: First ❶, to allow for hash-based comparisons between signals, the raw nanopore signals are processed and discretized robustly to signal noise and errors. Second ❷, overlapping regions between signals are detected by finding hash matches and using the Hough transform to detect matching diagonals. Third ❸, spurious matches are rejected by learned classifiers. Fourth❹, to tolerate small deviations between signals, the fuzzy de Bruijn graph is assembled using the overlap information by collapsing regions which differ slightly to a single node. Fifth❺, the final graph is produced by laying out contigs and performing additional graph cleaning steps to remove remaining errors. We provide the pseudocode of all steps in Supplementary Algorithms S1–S9.

**Figure 1:**
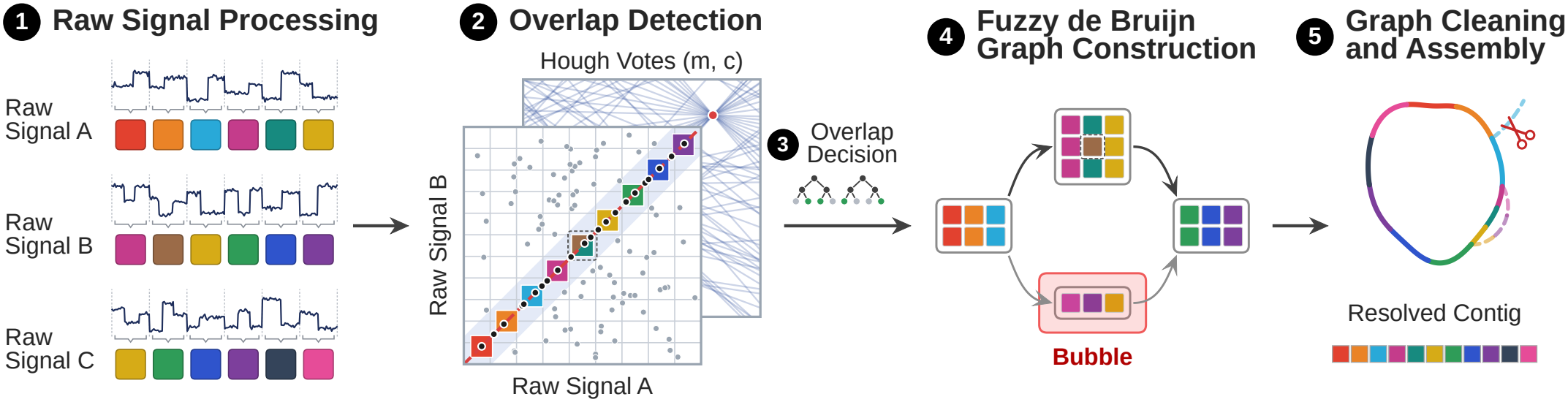
Overview of HARICOT.

### 2.1. Raw Signal Processing

The goal of the raw signal processing step is to convert each raw signal into a discrete signal sequence that can be compared across reads, so that matching regions between reads can later be found using hash-based seeds. During sequencing, as a molecule passes through the nanopore, a *k*-mer will occupy the pore and disrupt the ionic current. A segment in the signal corresponding to the occupancy of a *k*-mer is referred to as an *event*. When the molecule shifts and a new *k*-mer occupies the pore, this usually leads to a sharp change in current, delimiting the boundary of an event. However, the raw nanopore signal is significantly influenced by noise, which leads to fluctuations of signal values within events and varying event lengths.

To deal with these types of noise, HARICOT processes the raw signals in three steps, similar to prior raw signal analysis tools [8, 9, 18, 42]. First, to make different signals comparable, the raw nanopore signal is standardized and segmented into events using a statistical t-test [43, 44] that detects abrupt changes in current. Second, to reduce effects of noise within an event, each event is represented by a robust mean estimate and quantized to a discrete *bucket*. Third, to reduce oversegmentation, runs of identical buckets are joined, similar to how homopolymer compression is applied to nucleotide sequences.

While this sequence of buckets can in principle be treated like nucleotide sequences, e.g. by grouping *e* subsequent buckets to *e*-mers similar to *k*-mers, there are three main challenges which prevent the direct application for de Bruijn graph construction: 1) noise may shift the mean estimate per event, leading to different bucket assignments for the same underlying nucleotide sequence, 2) imprecise segmentation through the t-test, most commonly oversegmentation, and errors during sequencing such as skip errors lead to different sequences of buckets for the same underlying nucleotide sequence, and 3) due to quantization and merging of buckets, exact matches of e-mers are not specific, since two identical *e*-mers might be produced by very different underlying nucleotide sequences.

### 2.2 Overlap Detection

To form the nodes of the assembly graph from noisy seed matches between signals, HARICOT quickly and accurately identifies overlapping regions between signal pairs by utilizing the seed matches over bins with a Hough transform strategy in three steps.

**First**, to find the pairs of reads that may overlap, HARICOT stores the seeds of all reads in a hash table and collects the seed matches between each pair of reads, called *anchors*. An anchor (*p, p*^′^) gives the event positions of a shared seed in the two reads, where *p* is the position in the longer read. Figure 2 shows that the anchors of an overlap lie in a band around the line *p* = *mp*^′^ + *c* for an offset *c* and a slope *m* close to one. As in RawHash2 [18] and wtdbg2 [36], HARICOT masks the most frequent seeds, which removes uninformative anchors and reduces the runtime. To adapt to the sequencing depth and genome size, the masking threshold is a multiple of the mean seed frequency with a fixed lower bound. To avoid the costly second step for the many pairs of reads that share anchors only by chance, a *pre-filter* counts the anchors in wide bands of slope one, which contain the anchors of overlaps whose slopes differ slightly from one. The pre-filter keeps a pair only if its densest band contains significantly more anchors than expected for uniformly distributed anchors, using a lenient threshold to avoid discarding true overlaps.

**Figure 2:**
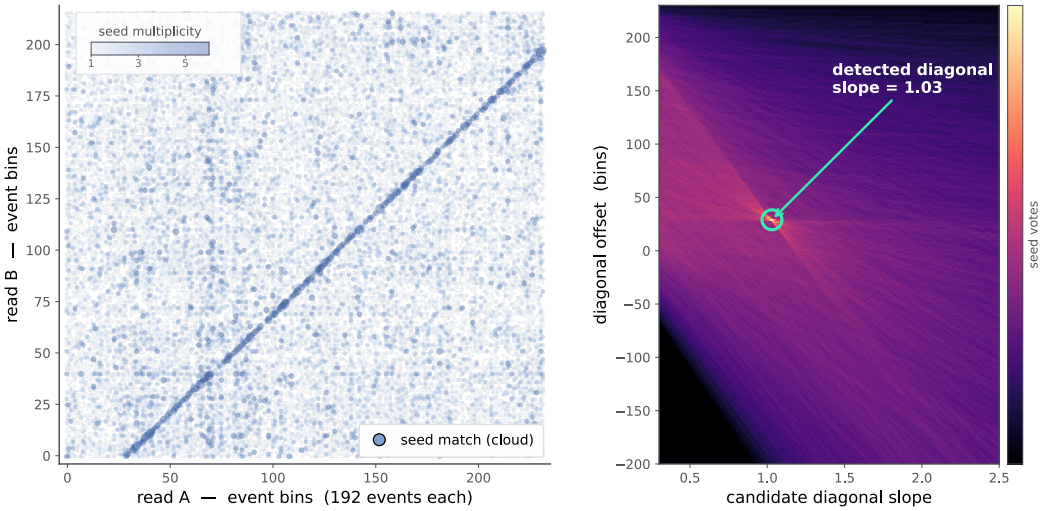
Seed matches over bins, with corresponding Hough transform.

**Second**, to find the line of an overlap among noisy anchors, HARICOT uses the Hough transform [38], a technique from computer vision to detect lines in images. This is because a single matching *e*-mer is weak evidence of an overlap, which makes the popular chaining strategy (i.e., connecting anchors with dynamic programming) of RawHash2 [18] ineffective when the goal is to find overlaps between noisy signals. Chaining fails since 1) many anchors are spurious, as Figure 2 shows, so chains can form from spurious anchors, and 2) noise removes all anchors in some regions of an overlap, so chains break at these gaps. The Hough transform avoids both problems by finding a line from all anchors close to it without connecting them into a chain. In the Hough transform, each anchor votes for the lines that pass through it. For a slope *m*, the line through an anchor has the offset *c* = *p −mp*^′^. The best line of slope *m* has the most votes for offsets in [*c −τ, c* +*τ*] (i.e., the most anchors within a distance *τ*). We call these anchors the *anchors of the line*. The tolerance *τ* accounts for noise in the anchor positions. To limit the number of tested slopes, HARICOT searches the slopes from coarse to fine ranges. A least-squares fit to the anchors of the best line refines its slope and offset.

To decide whether the line is a true overlap, HARICOT compares the number *n* of anchors of the line with the number expected by chance. By chance, *n* is approximately Poisson-distributed with mean *E*, the expected number of anchors of the line if the anchor positions of the two reads were paired at random. Comparing *n* directly with *E* is biased, since the largest of many counts exceeds their mean even by chance. The anchor offsets range from *c*_min_ to *c*_max_, which gives *M* = (*c*_max_ *− c*_min_)/2*τ* non-overlapping lines of slope *m*. Therefore, HARICOT computes

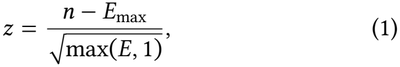

where *E*_max_, the (1 *−* 1/*M*)-quantile of the Poisson distribution, approximates the largest of *M* such counts. HARICOT discards the pairs with a low *z*. Since *M* grows with the read lengths, *E*_max_ keeps *z* comparable across read lengths.

**Third**, since the nodes of the graph are built from *bins* (i.e., groups of a fixed number of consecutive events), HARICOT converts the line into an *alignment path* between the bins of the two reads. The anchors of the line lie up to *τ* away from it, so the anchors in one bin of the longer read can fall into different bins of the other read. Therefore, HARICOT assigns each bin of the longer read to the bin of the other read that contains most of these anchors. Bins without anchors of the line remain *unassigned*. HARICOT discards the overlaps whose path is too short, contains too few matching events, or has too many unassigned bins, as we show in Supplementary Algorithm S2.

### 2.3. Overlap Decision

The goal of the overlap decision step is to remove the false overlaps that pass overlap detection while keeping as many true overlaps as possible. The detected overlaps contain false overlaps (e.g. due to repeats), and true overlaps whose path is misplaced. This is a crucial step, since a false overlap can merge two unrelated genomic regions into one node, while a missing overlap can break a contig.

Whether a detected overlap is true depends on several factors, such as the significance of its line, the repeat context of the two reads, and its consistency with the overlaps of other reads. Since it is unclear *a priori* how to combine these factors, HARICOT selects overlaps using two random forest classifiers [39], which combine the features of each overlap that we list in Supplementary Table S12. At contig ends, HARICOT uses a third classifier, as we explain in Section 2.5. To transfer the classifiers across genomes and depths, HARICOT normalizes each feature that grows with the depth, such as the number of overlapping reads, by its typical value in the same read or dataset.

The first classifier scores each overlap using four groups of features:

- **Alignment:** the significance, geometry and path shape of the overlap, e.g., its slope, overhangs and unassigned bins.
- **Repeat context:** the frequency of the shared seeds and the number of overlaps covering the region, relative to their typical values, since a repeat collects the seeds and reads of all its copies.
- **Consistency:** whether the common partners of the two reads overlap them at agreeing offsets.
- **Signal similarity:** the dynamic time warping (DTW) [45,46] cost between the unquantized event values of the two reads along the overlap, compared to an unrelated offset.

The second classifier targets the overlaps between repeat copies that the first classifier falsely accepts, which can be recognized in the context of the other accepted overlaps. For a true overlap between reads *A* and *B*, the reads that overlap *A* in the shared region should also overlap *B* at consistent offsets, whereas for an overlap between repeat copies these reads split into groups that do not overlap each other. The second classifier is therefore applied to the overlaps accepted by the first one and measures how many partners of *A* in the shared region also overlap *B* at consistent offsets, and vice versa, together with the first-stage scores of the overlap and of its neighbors.

We train the classifiers on datasets whose true overlaps are known from the positions of the reads on their reference genomes, as we describe in Section 3.1. True overlaps with a misplaced path are labeled as false, since they also harm the graph. Each decision threshold is set to keep a fixed fraction of the true training overlaps. Since overlaps are more redundant at higher depth, a stricter threshold is used for the second classifier at higher depth. The depth is estimated as the median number of reads covering a bin, using the overlaps kept at the most lenient threshold.

### 2.4. Fuzzy de Bruijn Graph Construction

The goal of graph construction is to collapse approximately equal *K*-bins of different reads that come from the same genomic region into a single node, using the selected overlaps.

The graph is constructed in four steps, the last three of which follow wtdbg2 [36]. **First**, since alignment paths are noisy and contain unassigned bins, which would break nodes, each path is straightened: a line is fitted to the path with the Theil–Sen estimator [47, 48], which is robust to misplaced bins. Then each bin is moved onto this line, and the unassigned bins are filled in. **Second**, each read is divided into *K*-bins, and each *K*-bin is projected onto the other read of each of its overlaps. If the projection spans between *K −* 1 and *K* + 1 bins, it is added to the candidate node of this *K*-bin. This allows for *fuzzy* matches between *K*-bins. **Third**, the candidates are resolved into a single tiling by committing them in decreasing order of supporting reads. Each committed node claims its intervals on the reads, blocking later candidates on the same region. Candidates supported by too many reads, which is typical of repeats, or left with too few unclaimed intervals are discarded. **Fourth**, nodes that appear consecutively on a read are connected by an edge.

### 2.5. Graph Cleaning and Assembly

The goal of graph cleaning is to remove errors caused by remaining false overlaps and by noise, so that long contigs can be laid out. The graph is simplified with the transitive reduction, tip trimming and bubble popping of wtdbg2 [36]. To better address the specific noise profile of raw signal graphs, HARICOT includes two additional cleaning steps. First, nodes with identical neighbors are merged. Such nodes are duplicates that two different reads create for the same region. Second, edges with low support relative to the coverage of neighboring nodes are removed, which adapts to the local depth. All cleaning steps are applied over several rounds, followed by tip trimming and bubble popping until the graph no longer changes.

To lay out contigs, HARICOT uses the fragment layout approach of wtdbg2. Since reads are usually much longer than branching regions, a read that spans a branching node shows how the graph continues across it. Therefore, the branching nodes are removed, which splits the graph into unbranched fragments, and fragments visited by the same read are linked. Weakly supported links, transitive links and links to short deadend fragments are removed, and the chains of uniquely linked fragments form the contigs.

Finally, HARICOT includes a *salvaging* procedure which joins contigs that are broken by a single missing overlap. Such an overlap might be missing because the classifiers falsely rejected it or because it was missed during overlap detection. The latter happens in particular when two longer reads only share a short overlap, which is then rejected as too insignificant.

We recover missing overlaps in two steps, considering only pairs of reads at the ends of different contigs. **First**, among the overlaps rejected by the classifiers, we consider those that connect two contig ends in a consistent orientation (i.e., overlaps from the right end of one contig to the left end of another, and containments). The best overlap at each contig end is the one with the highest first-stage score, and mutually best overlaps whose score exceeds a minimum are added. **Second**, to recover overlaps that detection missed, we test each line at a few slopes close to one against the number of anchors expected on this line by chance, instead of only the line with the most anchors, which favors long lines. A pair is kept if its most significant line whose anchors reach the read ends remains significant after correcting for the number of lines and pairs tested, and implies an overlap from the right end of one contig to the left end of another. Since overlap detection does not report the kept pairs as candidate overlaps, HARICOT scores the kept pairs using a third random forest classifier without the features of the detected line and path. HARICOT adds the accepted pairs that are the most significant at both contig ends.

## 3. Results

### 3.1. Evaluation Methodology

We implement HARICOT in Rust. HARICOT takes only raw signals from POD5 files as input and outputs its contigs in the Graphical Fragment Assembly (GFA) format, where each node of a contig is a range of events in one read, called the *anchor read* of the node.

We evaluate HARICOT in two parts, 1) the quality of the assemblies and 2) the computational resources, which we present in Sections 3.2 and 3.3. In both parts, we compare HARICOT with three assemblers. First, Rawsamble [20] is the only published raw signal assembler. Second, wtdbg2 [36] constructs the same type of assembly graph as HARICOT from basecalled reads. Third, Flye [40,41], a widely used assembler of nanopore reads, shows the contiguity that the basecalled reads of each dataset achieve. We basecall all reads with the super-accurate (SUP) model of Dorado [17].

#### Datasets

To cover genomes of different sizes and repeat content, we use 14 datasets, which we list in Table 1 and with their sources in Supplementary Table S3. D1–D6 are six chromosomes of five bacterial isolates [49] from SquiDBase [50]. D7–D9 are three chromosomes of *A. thaliana* from ENA [51]. D10–D14 come from ONT Open Data [52, 53]. D10– D13 are three chromosome arms and the X chromosome of *D. melanogaster*. D14 is chromosome 18 of the human genome HG002 with the reads of both haplotypes. We evaluate D14 against the maternal chromosome 18 of HG002 Q100 v1.2 [54].

**Table 1:** Datasets used in our evaluation.

| **Dataset** | **Organism (chromosome)** | **Length (Mb)** |
|---|---|---|
| D1 | *C. jejuni* | 1.8 |
| D2 | *L. monocytogenes* | 2.9 |
| D3, D4 | *V. parahaemolyticus* (1, 2) | 3.3, 1.9 |
| D5 | *S. enterica* | 4.8 |
| D6 | *S. dysgalactiae* | 2.2 |
| D7−D9 | *A. thaliana* (2, 3, 5) | 19.7−27.0 |
| D10−D13 | *D. melanogaster* (2L, 2R, 3L, X) | 23.5−28.1 |
| D14 | *H. sapiens* HG002 (18) | 78.9 |

Each dataset contains the reads of at least 5 kb whose basecalls align to its chromosome or chromosome arm, as we show in Supplementary Tables S4 and S5. Since HARICOT and Rawsamble cannot construct the reverse complement of a raw signal, we assemble each dataset from two types of read sets. First, the *forward-strand* read sets contain only the reads that align to the forward strand, at a *top depth* of at most 30× and at 12× and 6×. With these read sets, we compare the tools without the strand limitation and evaluate the effect of sequencing depth. Selecting these reads requires the reference genome. Second, the *both-strand* read sets contain the reads of both strands at the top depth, as in a real *de novo* assembly. We use the basecalls only to prepare the read sets, to label the training data and to evaluate the assemblies.

#### Classifier training

HARICOT uses three random forest classifiers to decide which overlaps to keep, as we explain in Section 2.3. To evaluate HARICOT on genomes that the classifiers do not see during training, we train the classifiers on 30 read sets of five other bacterial genomes [55, 56]. These genomes share no genus or sequencing run with the evaluation datasets, as Supplementary Table S3 shows. For each genome, we use six read sets: 1) the forward-strand reads at 30×, 12× and 6×, as for the evaluation datasets, and 2) fragments of 4 kb cut from the 30× reads, at the same three depths. We label each candidate overlap as true or false by aligning the base-called reads to their reference genomes with minimap2 [37]. We set the other parameters of HARICOT using the training genomes and four development datasets, which we do not use for evaluation, as we explain in Supplementary Section S3.

#### Assembly quality

We evaluate each assembly with QUAST [57, 58] against the reference of its dataset. Since HARICOT does not compute a consensus sequence and QUAST requires nucleotide sequences, we generate the sequence of each HARICOT contig from the basecalls of its anchor reads. To validate this approach, we generate the contig sequences of wtdbg2 in the same way and compare them with its consensus sequences in Supplementary Table S11. For Rawsamble, miniasm [35] generates the contig sequences from the basecalled reads. On 17 of the 56 read sets, miniasm cannot generate these sequences, since Rawsamble reports overlap positions in events rather than in bases. Rerunning miniasm on at most 25, 15 or 10 overlaps per read recovers 12 of these read sets, which we mark with † in the result tables.

We report various metrics in Tables 2 and 3 and Supplementary Tables S6–S11 to evaluate the contiguity, accuracy and completeness of the assemblies. First, the length of the longest contig and NG50 (i.e., the length of the shortest contig among the longest contigs that together reach half of the reference length) measure contiguity. We mark NG50 with a dash when it is undefined (i.e., when the contigs total less than half of the reference length). Second, the number of misassemblies (MA) measures accuracy. Third, the percentage of the reference that the contigs cover, called the genome fraction (GF), measures completeness. Since the contig sequences of HARICOT and Rawsamble are not consensus sequences, their lengths can differ slightly from those of the regions they cover. Therefore, when we mark the best and second best values in bold and underline in the result tables, values of the longest contig and NG50 that differ by less than 1% count as equal. Without consensus, the NG50 of wtdbg2 barely changes, whereas its genome fraction decreases and its number of misassemblies increases on most datasets, as Supplementary Table S11 shows. These results suggest that NG50 is comparable across all tools, while generating contig sequences from single reads likely lowers the genome fraction and raises the number of misassemblies of HARICOT and Rawsamble.

**Table 2:**
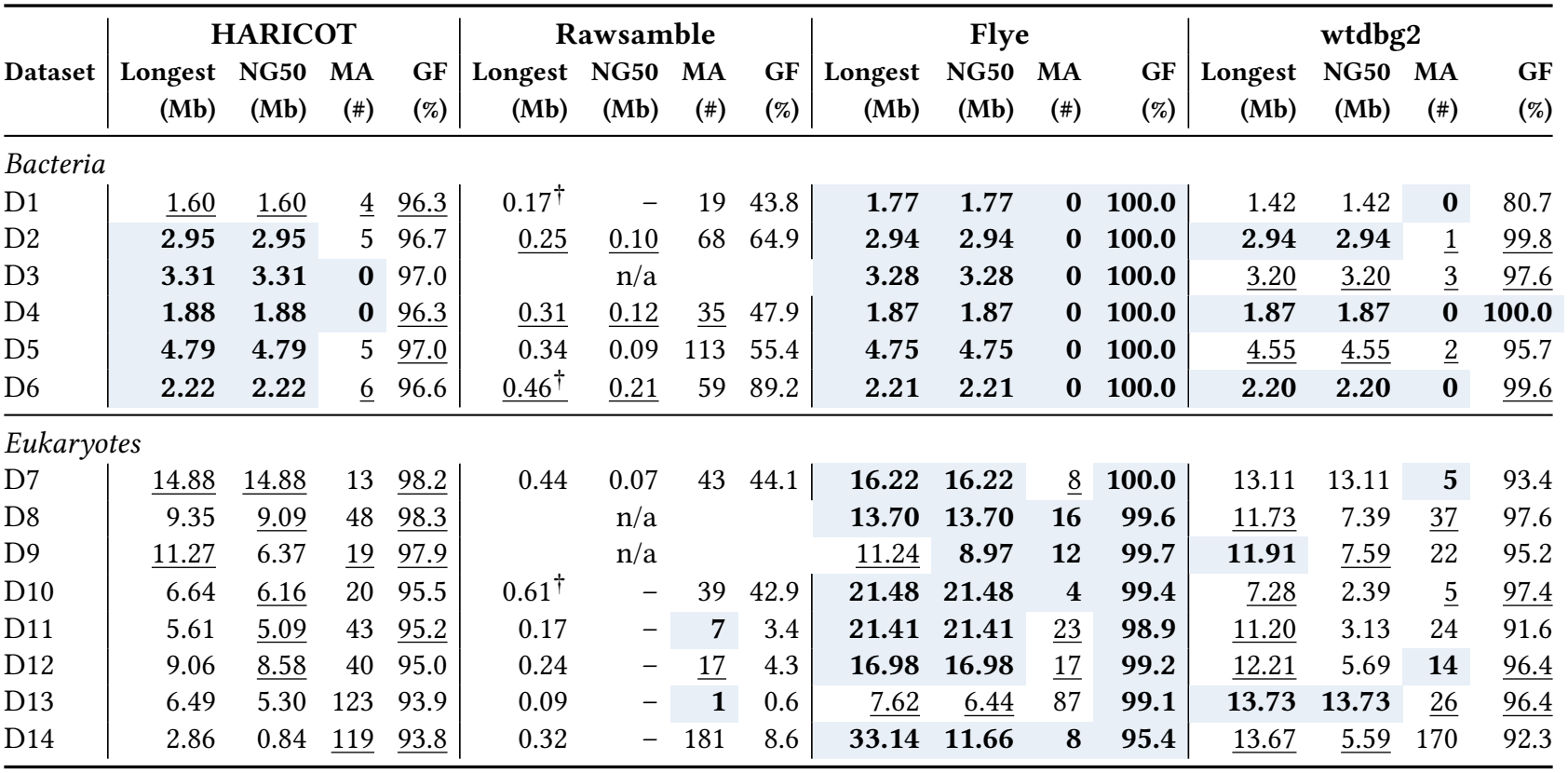
Assembly quality on the forward-strand datasets at the top depth. ^†^ Rawsamble retried by limiting overlaps per read. n/a: miniasm cannot generate the contig sequences of Rawsamble

|  | HARICOT |  |  |  | Rawsamble |  |  |  | Flye |  |  |  | wtdbg2 |  |  |  |
|---|---|---|---|---|---|---|---|---|---|---|---|---|---|---|---|---|
| Dataset | Longest | NG50 | MA | GF | Longest | NG50 | MA | GF | Longest | NG50 | MA | GF | Longest | NG50 | MA | GF |
|  | (Mb) | (Mb) | (#) | (%) | (Mb) | (Mb) | (#) | (%) | (Mb) | (Mb) | (#) | (%) | (Mb) | (Mb) | (#) | (%) |
| *Bacteria* |  |  |  |  |  |  |  |  |  |  |  |  |  |  |  |  |
| D1 | 1.60 | 1.60 | 4 | 96.3 | 0.17<sup>†</sup> | – | 19 | 43.8 | **1.77** | **1.77** | **0** | **100.0** | 1.42 | 1.42 | **0** | 80.7 |
| D2 | **2.95** | **2.95** | 5 | 96.7 | 0.25 | 0.10 | 68 | 64.9 | **2.94** | **2.94** | **0** | **100.0** | **2.94** | **2.94** | 1 | 99.8 |
| D3 | **3.31** | **3.31** | **0** | 97.0 | n/a |  |  |  | **3.28** | **3.28** | **0** | **100.0** | 3.20 | 3.20 | 3 | 97.6 |
| D4 | **1.88** | **1.88** | **0** | 96.3 | 0.31 | 0.12 | 35 | 47.9 | **1.87** | **1.87** | **0** | **100.0** | **1.87** | **1.87** | **0** | **100.0** |
| D5 | **4.79** | **4.79** | 5 | 97.0 | 0.34 | 0.09 | 113 | 55.4 | **4.75** | **4.75** | **0** | **100.0** | 4.55 | 4.55 | 2 | 95.7 |
| D6 | **2.22** | **2.22** | 6 | 96.6 | 0.46<sup>†</sup> | 0.21 | 59 | 89.2 | **2.21** | **2.21** | **0** | **100.0** | **2.20** | **2.20** | **0** | 99.6 |
| *Eukaryotes* |  |  |  |  |  |  |  |  |  |  |  |  |  |  |  |  |
| D7 | 14.88 | 14.88 | 13 | 98.2 | 0.44 | 0.07 | 43 | 44.1 | **16.22** | **16.22** | 8 | **100.0** | 13.11 | 13.11 | **5** | 93.4 |
| D8 | 9.35 | 9.09 | 48 | 98.3 | n/a |  |  |  | **13.70** | **13.70** | **16** | **99.6** | 11.73 | 7.39 | 37 | 97.6 |
| D9 | 11.27 | 6.37 | 19 | 97.9 | n/a |  |  |  | 11.24 | **8.97** | **12** | **99.7** | **11.91** | 7.59 | 22 | 95.2 |
| D10 | 6.64 | 6.16 | 20 | 95.5 | 0.61<sup>†</sup> | – | 39 | 42.9 | **21.48** | **21.48** | **4** | **99.4** | 7.28 | 2.39 | 5 | 97.4 |
| D11 | 5.61 | 5.09 | 43 | 95.2 | 0.17 | – | **7** | 3.4 | **21.41** | **21.41** | 23 | **98.9** | 11.20 | 3.13 | 24 | 91.6 |
| D12 | 9.06 | 8.58 | 40 | 95.0 | 0.24 | – | 17 | 4.3 | **16.98** | **16.98** | 17 | **99.2** | 12.21 | 5.69 | **14** | 96.4 |
| D13 | 6.49 | 5.30 | 123 | 93.9 | 0.09 | – | **1** | 0.6 | 7.62 | 6.44 | 87 | **99.1** | **13.73** | **13.73** | 26 | 96.4 |
| D14 | 2.86 | 0.84 | 119 | 93.8 | 0.32 | – | 181 | 8.6 | **33.14** | **11.66** | **8** | **95.4** | 13.67 | 5.59 | 170 | 92.3 |

**Table 3:**
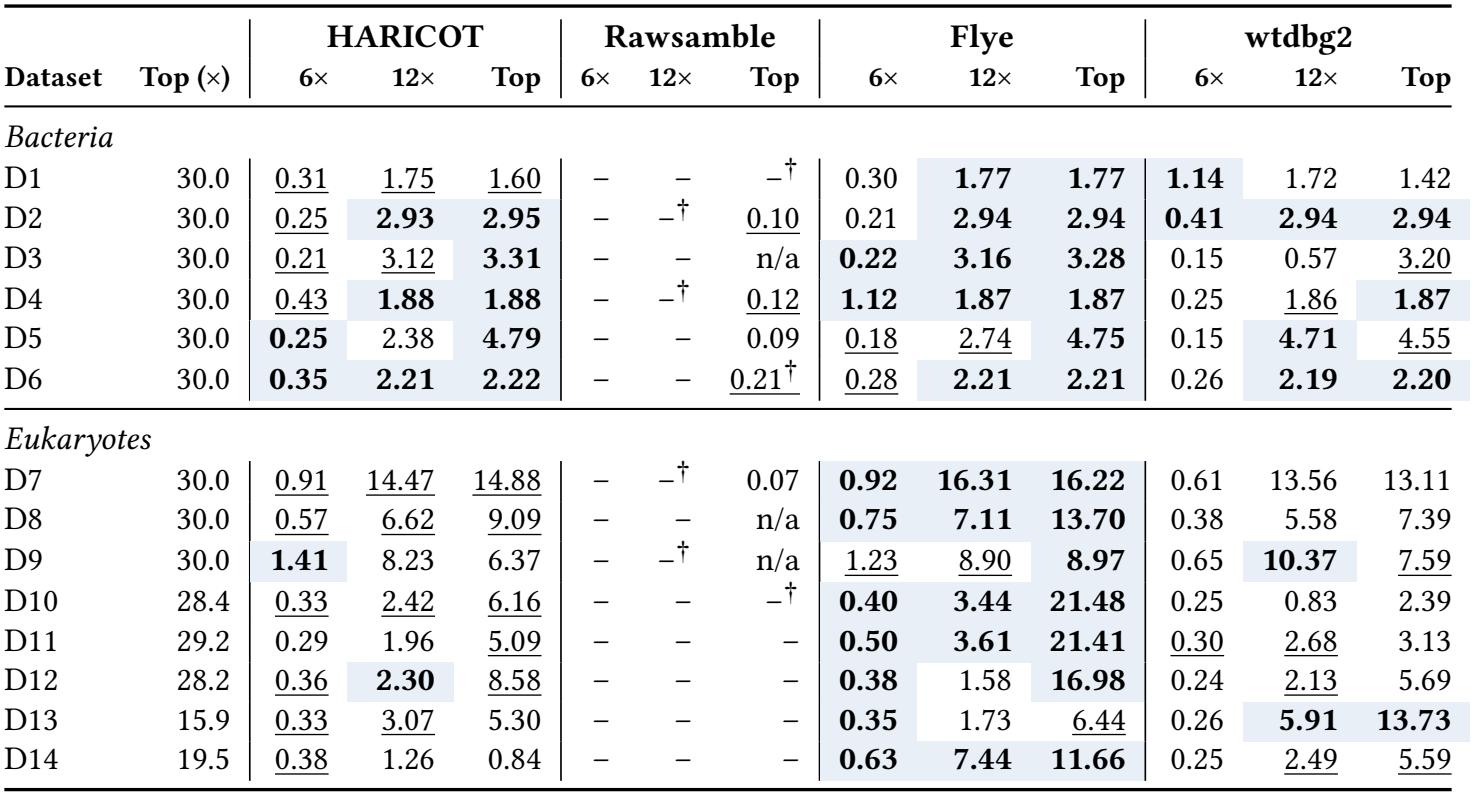
NG50 (Mb) at three sequencing depths. ^†^ Rawsamble retried by limiting overlaps per read. n/a: miniasm cannot generate the contig sequences of Rawsamble

|  |  | **HARICOT** |  |  | **Rawsamble** |  |  | **Flye** |  |  | **wtdbg2** |  |  |
|---|---|---|---|---|---|---|---|---|---|---|---|---|---|
| **Dataset** | **Top (×)** | **6×** | **12×** | **Top** | **6×** | **12×** | **Top** | **6×** | **12×** | **Top** | **6×** | **12×** | **Top** |
| *Bacteria* |  |  |  |  |  |  |  |  |  |  |  |  |  |
| D1 | 30.0 | 0.31 | 1.75 | 1.60 | – | – | –<sup>†</sup> | 0.30 | **1.77** | **1.77** | **1.14** | 1.72 | 1.42 |
| D2 | 30.0 | 0.25 | **2.93** | **2.95** | – | –<sup>†</sup> | 0.10 | 0.21 | **2.94** | **2.94** | **0.41** | **2.94** | **2.94** |
| D3 | 30.0 | 0.21 | 3.12 | **3.31** | – | – | n/a | **0.22** | **3.16** | **3.28** | 0.15 | 0.57 | 3.20 |
| D4 | 30.0 | 0.43 | **1.88** | **1.88** | – | –<sup>†</sup> | 0.12 | **1.12** | **1.87** | **1.87** | 0.25 | 1.86 | **1.87** |
| D5 | 30.0 | **0.25** | 2.38 | **4.79** | – | – | 0.09 | 0.18 | 2.74 | **4.75** | 0.15 | **4.71** | 4.55 |
| D6 | 30.0 | **0.35** | **2.21** | **2.22** | – | – | 0.21<sup>†</sup> | 0.28 | **2.21** | **2.21** | 0.26 | **2.19** | **2.20** |
| *Eukaryotes* |  |  |  |  |  |  |  |  |  |  |  |  |  |
| D7 | 30.0 | 0.91 | 14.47 | 14.88 | – | –<sup>†</sup> | 0.07 | **0.92** | **16.31** | **16.22** | 0.61 | 13.56 | 13.11 |
| D8 | 30.0 | 0.57 | 6.62 | 9.09 | – | – | n/a | **0.75** | **7.11** | **13.70** | 0.38 | 5.58 | 7.39 |
| D9 | 30.0 | **1.41** | 8.23 | 6.37 | – | –<sup>†</sup> | n/a | 1.23 | 8.90 | **8.97** | 0.65 | **10.37** | 7.59 |
| D10 | 28.4 | 0.33 | 2.42 | 6.16 | – | – | –<sup>†</sup> | **0.40** | **3.44** | **21.48** | 0.25 | 0.83 | 2.39 |
| D11 | 29.2 | 0.29 | 1.96 | 5.09 | – | – | – | **0.50** | **3.61** | **21.41** | 0.30 | 2.68 | 3.13 |
| D12 | 28.2 | 0.36 | **2.30** | 8.58 | – | – | – | **0.38** | 1.58 | **16.98** | 0.24 | 2.13 | 5.69 |
| D13 | 15.9 | 0.33 | 3.07 | 5.30 | – | – | – | **0.35** | 1.73 | 6.44 | 0.26 | **5.91** | **13.73** |
| D14 | 19.5 | 0.38 | 1.26 | 0.84 | – | – | – | **0.63** | **7.44** | **11.66** | 0.25 | 2.49 | 5.59 |

#### Performance

We run every tool with 16 threads on two Intel Xeon E5-2650 v4 CPUs, using the parameters and versions that we list in Supplementary Tables S1 and S2. For each tool, we measure the elapsed time, the CPU time and the peak memory usage of each step with /usr/bin/time and report the sum of the times and the largest peak memory over all steps. We report the elapsed time and the peak memory usage in Table 4 and all three measures in Supplementary Tables S7–S10. The times of Flye, wtdbg2 and miniasm do not include basecalling. In both parts, we report averages of ratios between tools as geometric means to reduce the effect of outliers.

**Table 4:**
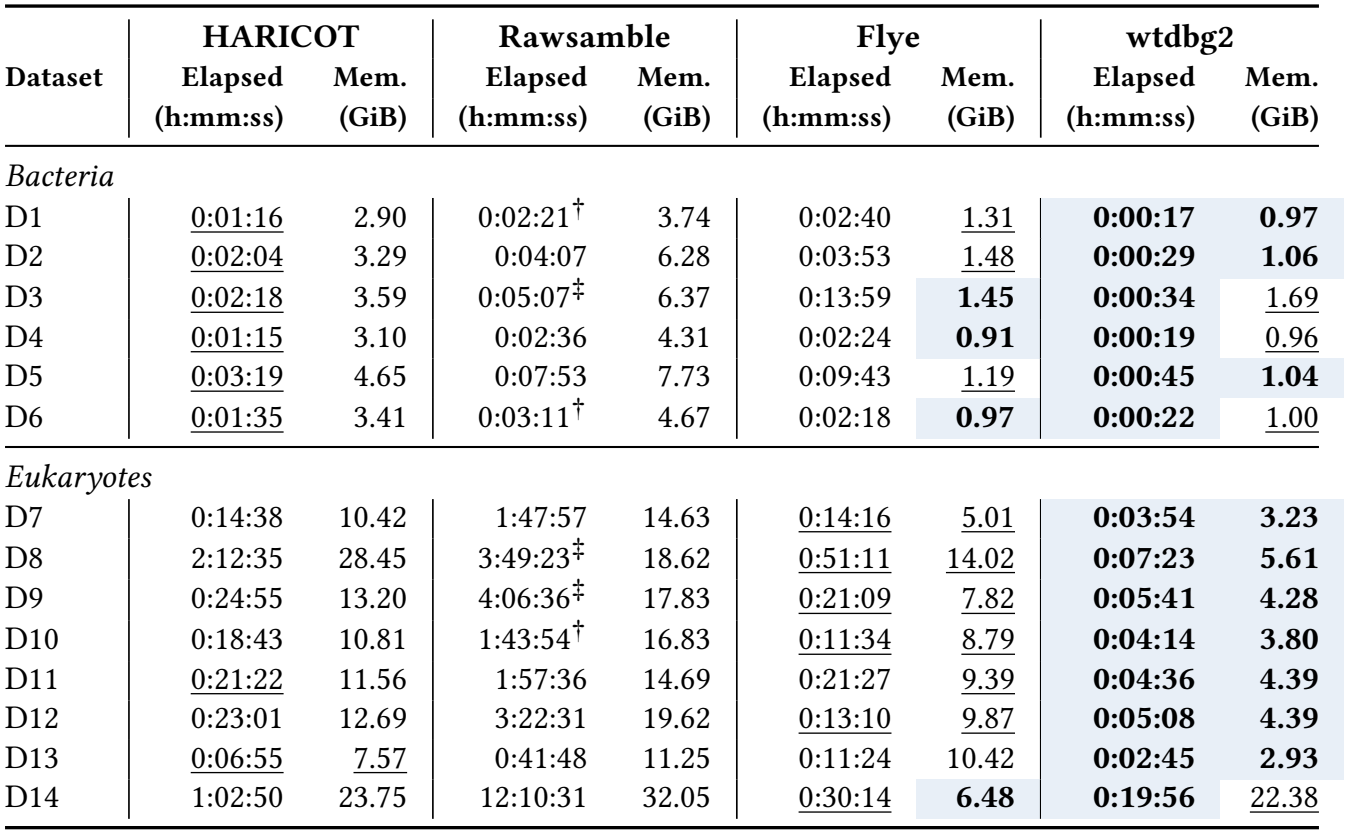
Elapsed time and peak memory. ^†^ Rawsamble retried by limiting overlaps per read. ^‡^ miniasm cannot generate the contig sequences of Rawsamble.

| **Dataset** | **HARICOT** |  | **Rawsamble** |  | **Flye** |  | **wtdbg2** |  |
|---|---|---|---|---|---|---|---|---|
|  | **Elapsed** | **Mem.** | **Elapsed** | **Mem.** | **Elapsed** | **Mem.** | **Elapsed** | **Mem.** |
|  | **(h:mm:ss)** | **(GiB)** | **(h:mm:ss)** | **(GiB)** | **(h:mm:ss)** | **(GiB)** | **(h:mm:ss)** | **(GiB)** |
| *Bacteria* |  |  |  |  |  |  |  |  |
| D1 | 0:01:16 | 2.90 | 0:02:21<sup>†</sup> | 3.74 | 0:02:40 | 1.31 | **0:00:17** | **0.97** |
| D2 | 0:02:04 | 3.29 | 0:04:07 | 6.28 | 0:03:53 | 1.48 | **0:00:29** | **1.06** |
| D3 | 0:02:18 | 3.59 | 0:05:07<sup>‡</sup> | 6.37 | 0:13:59 | **1.45** | **0:00:34** | 1.69 |
| D4 | 0:01:15 | 3.10 | 0:02:36 | 4.31 | 0:02:24 | **0.91** | **0:00:19** | 0.96 |
| D5 | 0:03:19 | 4.65 | 0:07:53 | 7.73 | 0:09:43 | 1.19 | **0:00:45** | **1.04** |
| D6 | 0:01:35 | 3.41 | 0:03:11<sup>†</sup> | 4.67 | 0:02:18 | **0.97** | **0:00:22** | 1.00 |
| *Eukaryotes* |  |  |  |  |  |  |  |  |
| D7 | 0:14:38 | 10.42 | 1:47:57 | 14.63 | 0:14:16 | 5.01 | **0:03:54** | **3.23** |
| D8 | 2:12:35 | 28.45 | 3:49:23<sup>‡</sup> | 18.62 | 0:51:11 | 14.02 | **0:07:23** | **5.61** |
| D9 | 0:24:55 | 13.20 | 4:06:36<sup>‡</sup> | 17.83 | 0:21:09 | 7.82 | **0:05:41** | **4.28** |
| D10 | 0:18:43 | 10.81 | 1:43:54<sup>†</sup> | 16.83 | 0:11:34 | 8.79 | **0:04:14** | **3.80** |
| D11 | 0:21:22 | 11.56 | 1:57:36 | 14.69 | 0:21:27 | 9.39 | **0:04:36** | **4.39** |
| D12 | 0:23:01 | 12.69 | 3:22:31 | 19.62 | 0:13:10 | 9.87 | **0:05:08** | **4.39** |
| D13 | 0:06:55 | 7.57 | 0:41:48 | 11.25 | 0:11:24 | 10.42 | **0:02:45** | **2.93** |
| D14 | 1:02:50 | 23.75 | 12:10:31 | 32.05 | 0:30:14 | **6.48** | **0:19:56** | 22.38 |

### 3.2. Assembly Quality

#### Contiguity and accuracy

Table 2 shows the assemblies of the forward-strand datasets at the top depth. Supplementary Figures S1 and S2 show the assembly graphs of all tools, which we draw with BandageNG, a fork of Bandage [59]. We make three key observations. First, HARICOT improves contiguity over Rawsamble on every dataset. HARICOT’s longest contigs are 5–72× longer than those of Rawsamble, and HARICOT covers 94–98% of each reference, whereas Rawsamble either fails or its contigs total less than half of the reference on 9 of the 14 datasets. This is because HARICOT tolerates noise both when it finds overlaps and when it constructs the graph, whereas Rawsamble passes its overlaps to miniasm, which is designed for the overlaps between basecalled reads. Second, on bacterial genomes, HARICOT produces near-complete assemblies. Its NG50 reaches the full chromosome length on five of the six datasets, matching Flye and equal to or higher than wtdbg2 on all six. HARICOT still makes a few misassemblies (0–6), while Flye assembles all bacterial datasets without any. Third, on eukaryotic genomes, HARICOT closes much of the gap to the basecalled assemblers but does not reach Flye. HARICOT achieves a higher NG50 than wtdbg2 on five of the seven *A. thaliana* and *D. melanogaster* datasets at a similar genome fraction, which shows that a fuzzy de Bruijn graph built directly from raw signals can match one built from basecalled sequences. Flye’s NG50 is 1.1–1.5× higher on *A. thaliana*, 1.2–4.2× higher on *D. melanogaster*, and about 14× higher on the human chromosome. HARICOT also makes more misassemblies than both basecalled assemblers on most eukaryotic datasets, likely due to false overlaps between repeat copies that remain after overlap selection (Section 4.1). We conclude that HARICOT assembles bacterial genomes as contiguously as basecalled assemblers and performs similarly to wtdbg2 on larger genomes, while more complex genomes remain challenging.

### Sequencing depth

Table 3 shows the NG50 of each tool at increasing depths of coverage. We make two key observations. First, HARICOT performs well at low and moderate depths. At 12×, it already produces near-complete assemblies for five of the six bacterial datasets, similar to Flye, and at 6×, its NG50 is higher than that of wtdbg2 on seven of the eight eukaryotic datasets. Rawsamble’s NG50 is undefined at both depths on all datasets. Second, the gap to Flye on eukaryotic genomes appears mainly at the top depth. At 12×, HARICOT’s NG50 is within about 10% of Flye on *A. thaliana* and higher than Flye on two *D. melanogaster* datasets (D12, D13). From 12× to the top depth, Flye’s NG50 on *D. melanogaster* grows by 4– 11×, while HARICOT’s grows by only 2–4×, and HARICOT’s NG50 even decreases on D1, D9 and D14. This suggests that HARICOT does not yet fully exploit additional reads, possibly because more reads also produce more false overlaps that overlap selection does not remove.

#### Both strands

Supplementary Table S6 shows the assemblies on datasets containing both strands. Since HARICOT cannot overlap reads from opposite strands, it assembles each strand separately, and its both-strand assemblies therefore contain each region of the genome once per strand. Using both strands increases HARICOT’s genome fraction to at least 98.9% on all non-human datasets, similar to Flye. Because each region appears twice, HARICOT’s NG50 and misassembly counts on these datasets are not directly comparable to those of Flye and wtdbg2, which merge both strands into a single assembly.

##### Key Result 1: HARICOT helps close the gap to basecalled assemblers

HARICOT produces near-complete bacterial assemblies comparable to Flye and, on larger eukaryotic genomes, achieves contiguity similar to wtdbg2 without basecalling.

### 3.3 Computational Resources

Table 4 shows the computational resources in terms of elapsed time and peak memory usage of all tools across the evaluated datasets. We make three key observations. First, HARICOT is faster than Rawsamble on all datasets, on average by 3.9× (up to 11.6×). The speedup grows from about 2× on the bacterial datasets to 5.5–11.6× on seven of the eight eukaryotic datasets, which shows that HARICOT scales better with genome size. HARICOT uses less memory than Rawsamble on all datasets except D8. Second, HARICOT’s runtime is comparable to that of Flye. HARICOT is faster than Flye on all bacterial datasets, and on eukaryotic datasets it ranges from 1.6× faster (D13) to 2.6× slower (D8). wtdbg2 is the fastest tool on all datasets. These runtimes exclude basecalling for Flye and wtdbg2, which HARICOT does not require. Third, HARICOT requires more memory than Flye and wtdbg2 on nearly all datasets, using 3–5 GiB on bacterial genomes compared to 1–2 GiB, and up to 28 GiB on D8. This is likely due to storing the seeds of all reads and the candidate overlaps between them during overlap detection. D8 is the most expensive dataset for HARICOT in both time and memory.

#### Key Result 2: HARICOT performs comparably to Flye at higher memory usage

HARICOT is faster than Rawsamble and comparable to Flye in runtime without requiring basecalling, at the cost of higher memory usage. wtdbg2 remains the fastest tool across all datasets.

## 4. Discussion and Future Work

### 4.1. Limitations

#### Overlap detection bottleneck

The requirement of computing pairwise similar regions between reads to find approximate matches between bins is the main computational bottleneck of HARICOT. In addition, errors in this step contribute directly to many of the downstream assembly errors, which need to be subsequently removed by additional filters and graph cleaning steps. It is possible that approximate matching could be further improved and accelerated using learned rules. For example, learned embeddings from raw signal windows encapsulating some contextual information could reduce the problem of approximate matching to finding similar embeddings. In addition, further improving the pre-filter step to remove false overlaps as early as possible could help further reduce unnecessary computations.

#### Reverse complements

A general weakness of current raw signal analysis tools, and therefore also of HARICOT, is their inability to deal with reverse complements. Some strategies have been developed to contend with this. For example, raw signal read mapping tools like RawHash [9, 18] build their indices using both forward and reverse strands of the reference. However, it is unclear how to apply these strategies to reference-free scenarios such as de novo assembly. Instead, HARICOT assembles strands separately, effectively halving the depth of coverage and leading to duplicate graphs. We leave the development of reverse-complementing in signal space as future work.

#### Short reads

Due to the binning and overlap detection strategy used in HARICOT, a certain minimum read length is required to assemble. While we use a minimum read length of 5 kb in our evaluations, it is possible that setting lower thresholds or reducing bin sizes may still yield similar quality and performance. However, HARICOT does not have an effective approach to make use of short reads without sacrificing robustness to noise. There is also trade-off in using shorter reads, as short nanopore reads tend to be of lower quality [60].

#### Structural read artifacts

HARICOT also deals poorly with structural read artifacts, such as chimeric reads, repeats and reads that contain both strands of the same region. While some effort has gone into making the overlap decision step robust to such artifacts, they nevertheless continue to lead to false overlaps. No dedicated step has been implemented to detect these artifacts, their detection from raw signals is left as future work.

### 4.2. Future Work

#### Streaming construction

To further accelerate assembly and make use of the real-time and selective sequencing capabilities of nanopore sequencers, it would be valuable to construct the assemblies in a streaming fashion as reads are sequenced. For nucleotide-space de Bruijn graphs, prior work Faucet [61] demonstrates streaming assembly of de Bruijn graphs, while Bifrost [62] explores better parallelization during construction. To apply this to raw signal assemblies, new algorithmic approaches are necessary, since several important steps of HARICOT, such as seed masking and depth-adaptive overlap selection, rely on statistics of the entire dataset.

#### Graph-based mapping

Several graph-based read mappers have been proposed for basecalled sequences [63–67], which have applications in particular for metagenomics [68, 69]. Recently, initial work in the raw signal space [23] has demonstrated graph mapping for raw signals as well. This opens the possibility of combining both raw signal assembly and raw signal mapping on graphs. For example, it might be possible to improve selective sequencing in metagenomic applications by mapping raw signals against the raw signal assembly to only selectively basecall certain reads.

#### Compression and de-duplication

The large size of raw signal data remains an obstacle to sharing datasets in the community and developing fast methods. Prior work in compressing raw nanopore signals has mostly focused on general compression techniques [70,71] that do not take biological structure and redundancy into account. Raw signal assemblies offer an opportunity to effectively detect redundancies in raw signals, which can be exploited to further improve compression.

#### Basecalling

The general goal of raw signal analysis is not to replace basecalling, but rather to augment and accelerate current analysis pipelines. We envision that raw signal assemblies can assist basecalling in moving from a linear sequence to a graph-based paradigm, wherein raw signals can be basecalled collectively to improve performance and reduce errors. For example, only one representative signal per node would need to be basecalled, or all signals of a node could be basecalled jointly to detect errors and modifications.

#### Consensus and assembly metrics

Since no existing raw signal assembler generates a consensus sequence, we translate raw signal assemblies back to nucleotide representations using a representative read to evaluate assembly quality. While this approach works to a certain extent, it is only a limited way of evaluating raw signal assemblies and can disadvantage raw signal assemblies in certain metrics such as NGA50. Developing methods that generate consensus sequences or signals directly from raw signals could enable a more holistic comparison with nucleotide assemblers. It is also necessary to develop new metrics tailored to raw signal assembly graphs.

## 5. Conclusion

We introduce HARICOT, the first raw signal assembler based on fuzzy de Bruijn graphs. HARICOT 1) identifies similar regions between signals using the Hough transform, 2) combines similar but not identical regions into nodes of the assembly graph, using statistical methods and learned classifiers to reject false overlaps, and 3) removes remaining errors with cleaning and salvaging steps adapted to the error profile of raw signal assemblies. Compared to Rawsamble, HARICOT constructs longest contigs that are on average 15.6× (up to 72×) longer while running on average 3.9× (up to 11.6×) faster. Without basecalling, HARICOT produces contigs spanning the entire chromosome on five of six bacterial chromosomes, and achieves a higher NG50 than wtdbg2 on five of seven plant and fly chromosomes, with a runtime comparable to Flye, which additionally requires basecalling. We believe that assembling raw signals before basecalling may open new directions, such as streaming assembly during sequencing and basecalling assembly graphs instead of individual reads.

## Supplementary Material for

### S1. Fuzzy de Bruijn graphs

De Bruijn graphs use short sequences of *k* subsequent nucleotides, or *k*-mers, as nodes in the assembly graph. Two nodes *a* and *b* in the assembly graph are connected by an edge if a (*k* + 1)-mer appears in a read, where *a* is the prefix and *b* the suffix of this (*k* + 1)-mer. The goal of the assembly is to reconstruct the underlying genome by finding a so-called *Eulerian* path through the graph which visits every edge once [1, 2]. In contrast to the problem of finding a path visiting every node in the graph as in the OLC paradigm, finding Eulerian paths is algorithmically efficient.

However, de Bruijn graphs come with trade-offs, the most important of which is their sensitivity to errors [3]. For instance, a single sequencing error will affect *k* different *k*-mers and lead to bubbles and divergences in the assembly graph. *Fuzzy* de Bruijn graphs, first introduced in wtdbg2 [4], address these limitations by allowing for inexact matches between items. To this end, longer sequences of nucleotides (on the order of tens or hundreds of nucleotides) are grouped into so-called *bins*. A node in the fuzzy de Bruijn graph then corresponds to K subsequent bins, or *K*-bins, and two nodes are connected by an edge if they appear subsequently on a read. To tolerate noise, K-bins which are “approximately equal” are collapsed to a single node, which differentiates fuzzy de Bruijn graphs from standard de Bruijn graphs requiring exact matches. To find approximately equal K-bins, we use information from overlaps between reads to find matching K-bins differing in at most one bin. Since overlaps are computed at the resolution of bins, fuzzy de Bruijn graphs are well suited to raw signals, where exact correspondences between individual positions of two signals are hard to find.

### S2. Configurations

#### S2.1. Parameters

Supplementary Table S1 lists the parameters of each tool.

**Table S1:**
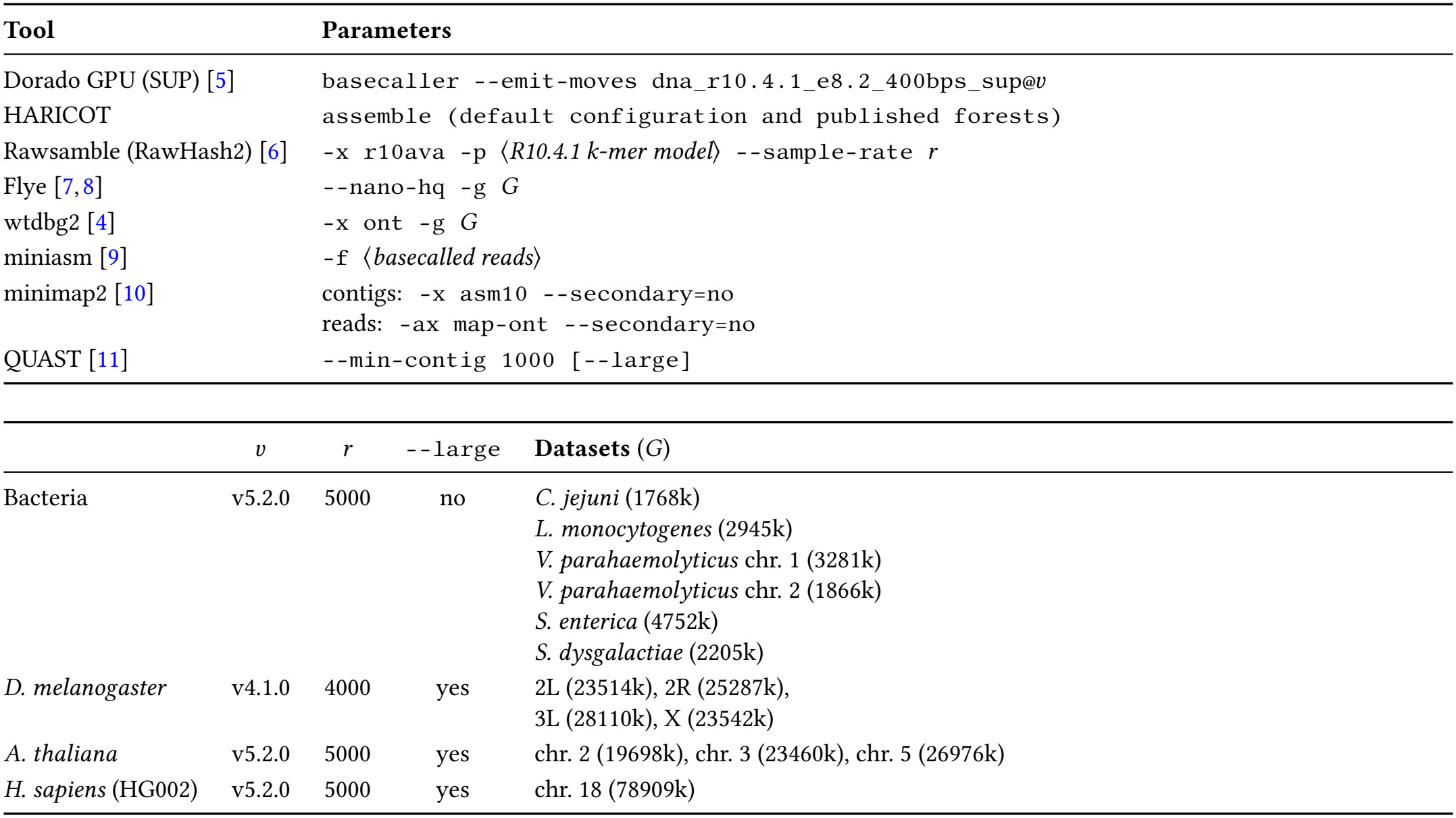
Parameters used in our evaluation. Top: command-line parameters per tool. Bottom: dataset-specific values substituted into those commands.

| Tool | Parameters |  |  |  |
|---|---|---|---|---|
| Dorado GPU (SUP) [5] | basecaller --emit-moves dna_r10.4.1_e8.2_400bps_sup@*v* |  |  |  |
| HARICOT | assemble (default configuration and published forests) |  |  |  |
| Rawsamble (RawHash2) [6] | -x r10ava -p ⟨*R10.4.1 k-mer model*⟩ --sample-rate *r* |  |  |  |
| Flye [7,8] | --nano-hq -g *G* |  |  |  |
| wtdbg2 [4] | -x ont -g *G* |  |  |  |
| miniasm [9] | -f ⟨*basecalled reads*⟩ |  |  |  |
| minimap2 [10] | contigs: -x asm10 --secondary=no<br>reads: -ax map-ont --secondary=no |  |  |  |
| QUAST [11] | --min-contig 1000 [--large] |  |  |  |
|  | *v* | *r* | --large | **Datasets** (*G*) |
| Bacteria | v5.2.0 | 5000 | no | *C. jejuni* (1768k)<br>*L. monocytogenes* (2945k)<br>*V. parahaemolyticus* chr. 1 (3281k)<br>*V. parahaemolyticus* chr. 2 (1866k)<br>*S. enterica* (4752k)<br>*S. dysgalactiae* (2205k) |
| *D. melanogaster* | v4.1.0 | 4000 | yes | 2L (23514k), 2R (25287k),<br>3L (28110k), X (23542k) |
| *A. thaliana* | v5.2.0 | 5000 | yes | chr. 2 (19698k), chr. 3 (23460k), chr. 5 (26976k) |
| *H. sapiens* (HG002) | v5.2.0 | 5000 | yes | chr. 18 (78909k) |

#### S2.2. Versions

Supplementary Table S2 lists the version of each tool. Scripts to reproduce all experiments can be found on https://github.com/STORMgroup/HARICOT. We use Dorado 1.4.0 [5] with --disable-read-splitting for the datasets sampled at 5 kHz, and Dorado 0.9.2, which cannot disable read splitting, for *D. melanogaster* (4 kHz).

**Table S2:** Versions of each tool and library.

| **Tool** | **Version** | **Link to the Source Code** |
|---|---|---|
| RawHash2 [12] | 2.5 | https://github.com/STORMgroup/RawHash2/tree/92366af |
| miniasm [9] | 0.3-r179 | https://github.com/lh3/miniasm/releases/tag/v0.3 |
| Flye [7,8] | 2.9.6-b1802 | https://github.com/mikolmogorov/Flye/releases/tag/2.9.6 |
| wtdbg2 [4] | 2.5 | https://github.com/ruanjue/wtdbg2/tree/b77c565 |
| minimap2 [10] | 2.31-r1302 | https://github.com/lh3/minimap2/releases/tag/v2.31 |
| QUAST [11] | 5.3.0 | https://github.com/ablab/quast/releases/tag/quast_5.3.0 |
| Dorado [5] | 0.9.2 | https://github.com/nanoporetech/dorado/releases/tag/v0.9.2 |
| Dorado [5] | 1.4.0 | https://github.com/nanoporetech/dorado/releases/tag/v1.4.0 |
| Bandage [13] | 2026.4.1 | https://github.com/asl/BandageNG/releases/tag/v2026.4.1 |

### S3. Datasets

Supplementary Table S3 lists all datasets used for evaluation (id starts with *D*) and training (id starts with *T*). We set the parameters of HARICOT using the training genomes and four development datasets, chromosomes 1 and 4 of *A. thaliana*, chromosome arm 3R of *D. melanogaster* and chromosome 20 of HG002, which we do not use for evaluation. These development datasets come from the same sequencing runs as D7–D14 but from different chromosomes.

**Table S3:** Datasets.

|  | Organism, strain | Reference | Length (Mb) | Device | Flow cells | Rate (kHz) | Source |
|---|---|---|---|---|---|---|---|
| D1 | *C. jejuni* ATCC 33560 | CP149499.1 | 1.77 | GridION | 4, multiplexed | 5 | SquiDBase SQB000011 [14] |
| D2 | *L. monocytogenes* EGD-e | CP149495.1 | 2.94 | GridION | 4, multiplexed | 5 | SquiDBase SQB000011 [14] |
| D3 | *V. parahaemolyticus* ATCC 17802 | CP149505.1 (chr. 1) | 3.28 | GridION | 4, multiplexed | 5 | SquiDBase SQB000011 [14] |
| D4 | *V. parahaemolyticus* ATCC 17802 | CP149506.1 (chr. 2) | 1.87 | GridION | 4, multiplexed | 5 | SquiDBase SQB000011 [14] |
| D5 | *S. enterica* ATCC 10708 | CP149507.1 | 4.75 | GridION | 4, multiplexed | 5 | SquiDBase SQB000011 [14] |
| D6 | *S. dysgalactiae* STG643 | CP149485.1 | 2.20 | GridION | FAX83388, multiplexed | 5 | SquiDBase SQB000011 [14] |
| D7 | *A. thaliana* Col-0 | NC_003071.7 (chr. 2) | 19.70 | P2 Solo | PBE26394 | 5 | ENA ERR17795792 [15] |
| D8 | *A. thaliana* Col-0 | NC_003074.8 (chr. 3) | 23.46 | P2 Solo | PBE26394 | 5 | ENA ERR17795792 [15] |
| D9 | *A. thaliana* Col-0 | NC_003076.8 (chr. 5) | 26.98 | P2 Solo | PBE26394 | 5 | ENA ERR17795792 [15] |
| D10 | *D. melanogaster* iso-1 | NT_033779.5 (2L) | 23.51 | MinION | FAV70669, FAV81161 | 4 | ONT Open Data [16] |
| D11 | *D. melanogaster* iso-1 | NT_033778.4 (2R) | 25.29 | MinION | FAV70669, FAV81161 | 4 | ONT Open Data [16] |
| D12 | *D. melanogaster* iso-1 | NT_037436.4 (3L) | 28.11 | MinION | FAV70669, FAV81161 | 4 | ONT Open Data [16] |
| D13 | *D. melanogaster* iso-1 | NC_004354.4 (X) | 23.54 | MinION | FAV70669, FAV81161 | 4 | ONT Open Data [16] |
| D14 | *H. sapiens* HG002 | chr18_MATERNAL [17] | 78.91 | PromethION | PAO89685 | 5 | ONT Open Data [18] |
| T1 | *E. coli* K-12 MG1655 | NC_000913.3 | 4.64 | MinION Mk1C | FAW79843, multiplexed | 5 | AWS Open Data [19] |
| T2 | *S. aureus* NO-MISS sample 8 | sample_8_2 contig_1 | 2.73 | P2 Integrated | PBK98658, multiplexed | 5 | ONT Open Data [20] |
| T3 | *H. pylori* 26695 | NC_018939.1 | 1.67 | MinION Mk1C | FAW83092 | 5 | AWS Open Data [19] |
| T4 | *T. denticola* ATCC 35405 | NC_002967.9 | 2.84 | PromethION | PBC50044, multiplexed | 5 | AWS Open Data [19] |
| T5 | *Nostoc* sp. PCC 7120 | NC_003272.1 | 6.41 | PromethION | PBC50044, multiplexed | 5 | AWS Open Data [19] |

Supplementary Table S4 lists the steps applied to select reads for training and evaluation, with Supplementary Table S5 showing the results of the selection on read counts.

**Table S4:** Read selection steps.

|  | Step | Rule | Datasets |
|---|---|---|---|
| 1 | Assignment | Keep reads whose primary alignment (minimap2 -ax map-ont) is on the chromosome with MAPQ ≥ 10. Bacteria: against a panel of the 14 assemblies of [21] and the five training references (for T1 and T2, without the other assembly of their species). D14: against the maternal haplotype, then on either chromosome 18 of the diploid reference. | all |
| 2 | Cross-species | Remove reads with an alignment of ≥ 1 kb to another species of the panel. | D1–D6, T1–T5 |
| 3 | Read length | Basecall ≥ 5 kb. | all |
| 4 | Strand | Strand of the primary alignment. | all |
| 5 | Top depth | Per strand, reads in the order of the MD5 hash of their ids up to 30×, or all reads. | all |
| 6 | Lower depths | 12× and 6× forward-strand subsets of the top set (6× ⊂ 12× ⊂ top). | all |
| 7 | Both strands | Forward and reverse top sets together. | D1–D14 |

**Table S5:** Reads at each read selection step.

|  |  |  |  | Eligible |  | Read sets |  |  |  | Top |
|---|---|---|---|---|---|---|---|---|---|---|
| **Dataset** | **Assigned** | **Cross-species** | **< 5 kb** | **Forward** | **Reverse** | **Top** | **12×** | **6×** | **Both** | **depth (×)** |
| D1 | 49,238 | 168 | 19,332 | 14,980 | 14,758 | 4,850 | 1,955 | 1,018 | 9,672 | 30.0 |
| D2 | 91,258 | 304 | 41,459 | 24,762 | 24,733 | 8,308 | 3,315 | 1,675 | 16,555 | 30.0 |
| D3 | 112,948 | 553 | 63,485 | 24,698 | 24,212 | 9,628 | 3,877 | 1,938 | 19,299 | 30.0 |
| D4 | 57,177 | 250 | 31,707 | 12,601 | 12,619 | 5,428 | 2,113 | 1,082 | 10,881 | 30.0 |
| D5 | 173,374 | 1,163 | 99,762 | 36,206 | 36,243 | 14,711 | 5,832 | 2,940 | 29,492 | 30.0 |
| D6 | 63,249 | 28 | 26,786 | 18,060 | 18,375 | 5,566 | 2,246 | 1,111 | 11,119 | 30.0 |
| D7 | 63,314 | – | 4,205 | 29,739 | 29,370 | 19,488 | 7,712 | 3,880 | 39,262 | 30.0 |
| D8 | 87,355 | – | 5,935 | 40,643 | 40,777 | 23,837 | 9,489 | 4,730 | 47,593 | 30.0 |
| D9 | 86,525 | – | 5,752 | 40,219 | 40,554 | 26,823 | 10,793 | 5,408 | 53,634 | 30.0 |
| D10 | 129,294 | – | 48,426 | 40,194 | 40,674 | 40,194 | 17,171 | 8,541 | 80,868 | 28.4 |
| D11 | 143,156 | – | 53,610 | 44,813 | 44,733 | 44,813 | 18,462 | 9,262 | 89,546 | 29.2 |
| D12 | 151,694 | – | 56,688 | 47,472 | 47,534 | 47,472 | 20,222 | 10,101 | 95,006 | 28.2 |
| D13 | 77,923 | – | 30,739 | 23,511 | 23,673 | 23,511 | 17,729 | 8,988 | 47,184 | 15.9 |
| D14 | 125,729 | – | 11,372 | 57,149 | 57,208 | 57,149 | 35,199 | 17,515 | 114,357 | 19.5 |
| T1 | 39,568 | 1 | 10,342 | 14,185 | 15,040 | 8,962 | 3,645 | 1,834 | – | 30.0 |
| T2 | 109,617 | 13 | 62,578 | 23,491 | 23,535 | 10,161 | 4,185 | 2,047 | – | 30.0 |
| T3 | 17,005 | 0 | 7,703 | 4,473 | 4,829 | 4,412 | 1,825 | 856 | – | 30.0 |
| T4 | 22,003 | 166 | 1,916 | 10,012 | 9,909 | 6,279 | 2,528 | 1,236 | – | 30.0 |
| T5 | 43,495 | 277 | 4,246 | 19,697 | 19,275 | 15,932 | 6,307 | 3,102 | – | 30.0 |

### S4. Extended Results

Supplementary Table S6 shows the results of running the assemblies on datasets containing both strands. Since the raw signal assemblers Rawsamble [6] and HARICOT do not support reverse complementing, strands will be assembled separately.

**Table S6:**
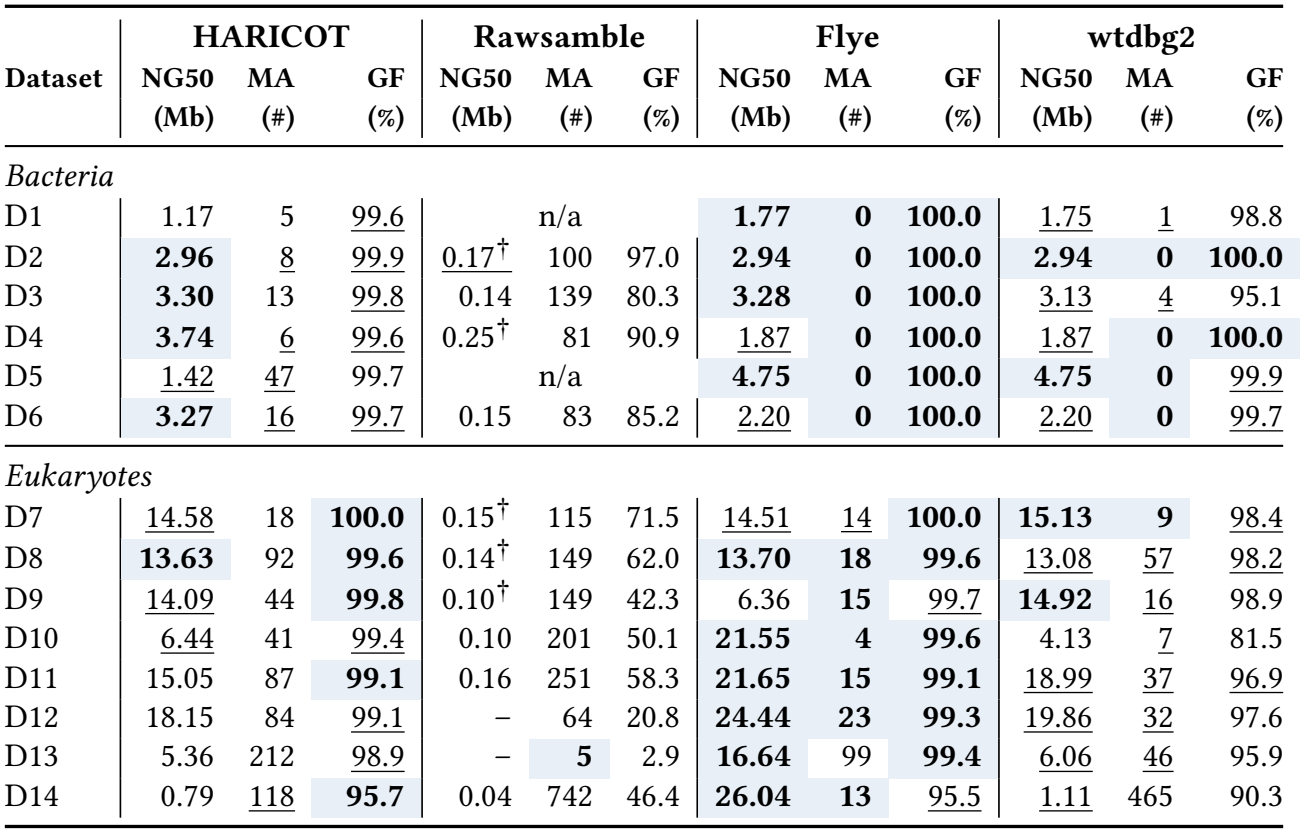
Assembly quality on the both-strand datasets. † Rawsamble retried by limiting overlaps per read. n/a: miniasm cannot generate the contig sequences of Rawsamble.

|  | HARICOT |  |  | Rawsamble |  |  | Flye |  |  | wtdbg2 |  |  |
|---|---|---|---|---|---|---|---|---|---|---|---|---|
| Dataset | NG50<br>(Mb) | MA<br>(#) | GF<br>(%) | NG50<br>(Mb) | MA<br>(#) | GF<br>(%) | NG50<br>(Mb) | MA<br>(#) | GF<br>(%) | NG50<br>(Mb) | MA<br>(#) | GF<br>(%) |
| *Bacteria* |  |  |  |  |  |  |  |  |  |  |  |  |
| D1 | 1.17 | 5 | 99.6 | n/a |  |  | **1.77** | **0** | **100.0** | 1.75 | 1 | 98.8 |
| D2 | **2.96** | 8 | 99.9 | 0.17<sup>†</sup> | 100 | 97.0 | **2.94** | **0** | **100.0** | **2.94** | **0** | **100.0** |
| D3 | **3.30** | 13 | 99.8 | 0.14 | 139 | 80.3 | **3.28** | **0** | **100.0** | 3.13 | 4 | 95.1 |
| D4 | **3.74** | 6 | 99.6 | 0.25<sup>†</sup> | 81 | 90.9 | 1.87 | **0** | **100.0** | 1.87 | **0** | **100.0** |
| D5 | 1.42 | 47 | 99.7 | n/a |  |  | **4.75** | **0** | **100.0** | **4.75** | **0** | 99.9 |
| D6 | **3.27** | 16 | 99.7 | 0.15 | 83 | 85.2 | 2.20 | **0** | **100.0** | 2.20 | **0** | 99.7 |
| *Eukaryotes* |  |  |  |  |  |  |  |  |  |  |  |  |
| D7 | 14.58 | 18 | **100.0** | 0.15<sup>†</sup> | 115 | 71.5 | 14.51 | 14 | **100.0** | **15.13** | **9** | 98.4 |
| D8 | **13.63** | 92 | **99.6** | 0.14<sup>†</sup> | 149 | 62.0 | **13.70** | **18** | **99.6** | 13.08 | 57 | 98.2 |
| D9 | 14.09 | 44 | **99.8** | 0.10<sup>†</sup> | 149 | 42.3 | 6.36 | **15** | 99.7 | **14.92** | 16 | 98.9 |
| D10 | 6.44 | 41 | 99.4 | 0.10 | 201 | 50.1 | **21.55** | **4** | **99.6** | 4.13 | 7 | 81.5 |
| D11 | 15.05 | 87 | **99.1** | 0.16 | 251 | 58.3 | **21.65** | **15** | **99.1** | 18.99 | 37 | 96.9 |
| D12 | 18.15 | 84 | 99.1 | – | 64 | 20.8 | **24.44** | **23** | **99.3** | 19.86 | 32 | 97.6 |
| D13 | 5.36 | 212 | 98.9 | – | **5** | 2.9 | **16.64** | 99 | **99.4** | 6.06 | 46 | 95.9 |
| D14 | 0.79 | 118 | **95.7** | 0.04 | 742 | 46.4 | **26.04** | **13** | 95.5 | 1.11 | 465 | 90.3 |

In Supplementary Tables S7, S8, S9, S10 we give extended results of assembly quality and computational resources for HARICOT, Rawsamble [6], Flye [7, 8], and wtdbg2 [4] respectively.

**Table S7:**
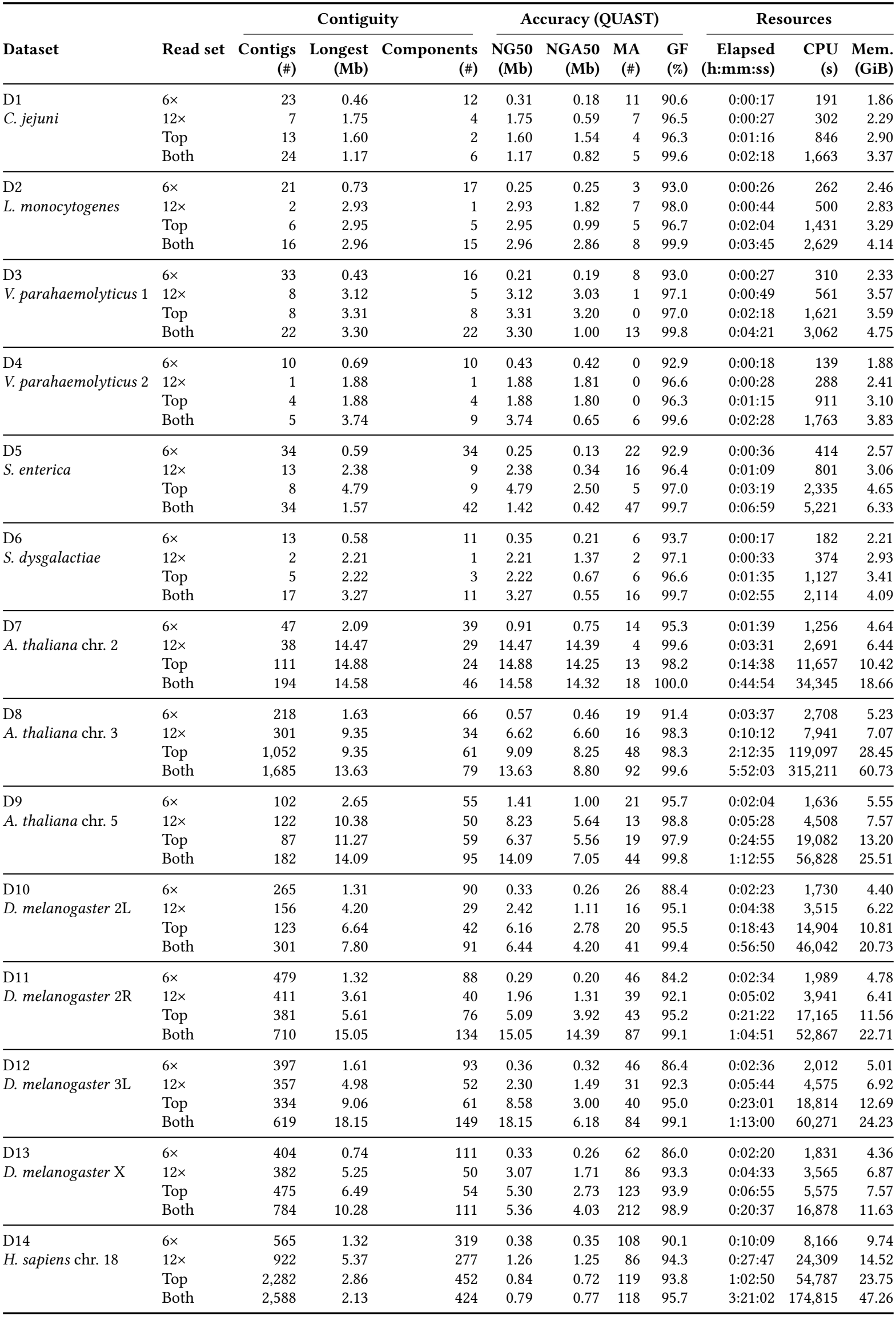
Assemblies by HARICOT.

|  |  | Contiguity |  |  | Accuracy (QUAST) |  |  |  | Resources |  |  |
|---|---|---|---|---|---|---|---|---|---|---|---|
| Dataset | Read set | Contigs<br>(#) | Longest<br>(Mb) | Components<br>(#) | NG50<br>(Mb) | NGA50<br>(Mb) | MA<br>(#) | GF<br>(%) | Elapsed<br>(h:mm:ss) | CPU<br>(s) | Mem.<br>(GiB) |
| D1 | 6× | 23 | 0.46 | 12 | 0.31 | 0.18 | 11 | 90.6 | 0:00:17 | 191 | 1.86 |
| *C. jejuni* | 12× | 7 | 1.75 | 4 | 1.75 | 0.59 | 7 | 96.5 | 0:00:27 | 302 | 2.29 |
|  | Top | 13 | 1.60 | 2 | 1.60 | 1.54 | 4 | 96.3 | 0:01:16 | 846 | 2.90 |
|  | Both | 24 | 1.17 | 6 | 1.17 | 0.82 | 5 | 99.6 | 0:02:18 | 1,663 | 3.37 |
| D2 | 6× | 21 | 0.73 | 17 | 0.25 | 0.25 | 3 | 93.0 | 0:00:26 | 262 | 2.46 |
| *L. monocytogenes* | 12× | 2 | 2.93 | 1 | 2.93 | 1.82 | 7 | 98.0 | 0:00:44 | 500 | 2.83 |
|  | Top | 6 | 2.95 | 5 | 2.95 | 0.99 | 5 | 96.7 | 0:02:04 | 1,431 | 3.29 |
|  | Both | 16 | 2.96 | 15 | 2.96 | 2.86 | 8 | 99.9 | 0:03:45 | 2,629 | 4.14 |
| D3 | 6× | 33 | 0.43 | 16 | 0.21 | 0.19 | 8 | 93.0 | 0:00:27 | 310 | 2.33 |
| *V. parahaemolyticus* 1 | 12× | 8 | 3.12 | 5 | 3.12 | 3.03 | 1 | 97.1 | 0:00:49 | 561 | 3.57 |
|  | Top | 8 | 3.31 | 8 | 3.31 | 3.20 | 0 | 97.0 | 0:02:18 | 1,621 | 3.59 |
|  | Both | 22 | 3.30 | 22 | 3.30 | 1.00 | 13 | 99.8 | 0:04:21 | 3,062 | 4.75 |
| D4 | 6× | 10 | 0.69 | 10 | 0.43 | 0.42 | 0 | 92.9 | 0:00:18 | 139 | 1.88 |
| *V. parahaemolyticus* 2 | 12× | 1 | 1.88 | 1 | 1.88 | 1.81 | 0 | 96.6 | 0:00:28 | 288 | 2.41 |
|  | Top | 4 | 1.88 | 4 | 1.88 | 1.80 | 0 | 96.3 | 0:01:15 | 911 | 3.10 |
|  | Both | 5 | 3.74 | 9 | 3.74 | 0.65 | 6 | 99.6 | 0:02:28 | 1,763 | 3.83 |
| D5 | 6× | 34 | 0.59 | 34 | 0.25 | 0.13 | 22 | 92.9 | 0:00:36 | 414 | 2.57 |
| *S. enterica* | 12× | 13 | 2.38 | 9 | 2.38 | 0.34 | 16 | 96.4 | 0:01:09 | 801 | 3.06 |
|  | Top | 8 | 4.79 | 9 | 4.79 | 2.50 | 5 | 97.0 | 0:03:19 | 2,335 | 4.65 |
|  | Both | 34 | 1.57 | 42 | 1.42 | 0.42 | 47 | 99.7 | 0:06:59 | 5,221 | 6.33 |
| D6 | 6× | 13 | 0.58 | 11 | 0.35 | 0.21 | 6 | 93.7 | 0:00:17 | 182 | 2.21 |
| *S. dysgalactiae* | 12× | 2 | 2.21 | 1 | 2.21 | 1.37 | 2 | 97.1 | 0:00:33 | 374 | 2.93 |
|  | Top | 5 | 2.22 | 3 | 2.22 | 0.67 | 6 | 96.6 | 0:01:35 | 1,127 | 3.41 |
|  | Both | 17 | 3.27 | 11 | 3.27 | 0.55 | 16 | 99.7 | 0:02:55 | 2,114 | 4.09 |
| D7 | 6× | 47 | 2.09 | 39 | 0.91 | 0.75 | 14 | 95.3 | 0:01:39 | 1,256 | 4.64 |
| *A. thaliana* chr. 2 | 12× | 38 | 14.47 | 29 | 14.47 | 14.39 | 4 | 99.6 | 0:03:31 | 2,691 | 6.44 |
|  | Top | 111 | 14.88 | 24 | 14.88 | 14.25 | 13 | 98.2 | 0:14:38 | 11,657 | 10.42 |
|  | Both | 194 | 14.58 | 46 | 14.58 | 14.32 | 18 | 100.0 | 0:44:54 | 34,345 | 18.66 |
| D8 | 6× | 218 | 1.63 | 66 | 0.57 | 0.46 | 19 | 91.4 | 0:03:37 | 2,708 | 5.23 |
| *A. thaliana* chr. 3 | 12× | 301 | 9.35 | 34 | 6.62 | 6.60 | 16 | 98.3 | 0:10:12 | 7,941 | 7.07 |
|  | Top | 1,052 | 9.35 | 61 | 9.09 | 8.25 | 48 | 98.3 | 2:12:35 | 119,097 | 28.45 |
|  | Both | 1,685 | 13.63 | 79 | 13.63 | 8.80 | 92 | 99.6 | 5:52:03 | 315,211 | 60.73 |
| D9 | 6× | 102 | 2.65 | 55 | 1.41 | 1.00 | 21 | 95.7 | 0:02:04 | 1,636 | 5.55 |
| *A. thaliana* chr. 5 | 12× | 122 | 10.38 | 50 | 8.23 | 5.64 | 13 | 98.8 | 0:05:28 | 4,508 | 7.57 |
|  | Top | 87 | 11.27 | 59 | 6.37 | 5.56 | 19 | 97.9 | 0:24:55 | 19,082 | 13.20 |
|  | Both | 182 | 14.09 | 95 | 14.09 | 7.05 | 44 | 99.8 | 1:12:55 | 56,828 | 25.51 |
| D10 | 6× | 265 | 1.31 | 90 | 0.33 | 0.26 | 26 | 88.4 | 0:02:23 | 1,730 | 4.40 |
| *D. melanogaster* 2L | 12× | 156 | 4.20 | 29 | 2.42 | 1.11 | 16 | 95.1 | 0:04:38 | 3,515 | 6.22 |
|  | Top | 123 | 6.64 | 42 | 6.16 | 2.78 | 20 | 95.5 | 0:18:43 | 14,904 | 10.81 |
|  | Both | 301 | 7.80 | 91 | 6.44 | 4.20 | 41 | 99.4 | 0:56:50 | 46,042 | 20.73 |
| D11 | 6× | 479 | 1.32 | 88 | 0.29 | 0.20 | 46 | 84.2 | 0:02:34 | 1,989 | 4.78 |
| *D. melanogaster* 2R | 12× | 411 | 3.61 | 40 | 1.96 | 1.31 | 39 | 92.1 | 0:05:02 | 3,941 | 6.41 |
|  | Top | 381 | 5.61 | 76 | 5.09 | 3.92 | 43 | 95.2 | 0:21:22 | 17,165 | 11.56 |
|  | Both | 710 | 15.05 | 134 | 15.05 | 14.39 | 87 | 99.1 | 1:04:51 | 52,867 | 22.71 |
| D12 | 6× | 397 | 1.61 | 93 | 0.36 | 0.32 | 46 | 86.4 | 0:02:36 | 2,012 | 5.01 |
| *D. melanogaster* 3L | 12× | 357 | 4.98 | 52 | 2.30 | 1.49 | 31 | 92.3 | 0:05:44 | 4,575 | 6.92 |
|  | Top | 334 | 9.06 | 61 | 8.58 | 3.00 | 40 | 95.0 | 0:23:01 | 18,814 | 12.69 |
|  | Both | 619 | 18.15 | 149 | 18.15 | 6.18 | 84 | 99.1 | 1:13:00 | 60,271 | 24.23 |
| D13 | 6× | 404 | 0.74 | 111 | 0.33 | 0.26 | 62 | 86.0 | 0:02:20 | 1,831 | 4.36 |
| *D. melanogaster* X | 12× | 382 | 5.25 | 50 | 3.07 | 1.71 | 86 | 93.3 | 0:04:33 | 3,565 | 6.87 |
|  | Top | 475 | 6.49 | 54 | 5.30 | 2.73 | 123 | 93.9 | 0:06:55 | 5,575 | 7.57 |
|  | Both | 784 | 10.28 | 111 | 5.36 | 4.03 | 212 | 98.9 | 0:20:37 | 16,878 | 11.63 |
| D14 | 6× | 565 | 1.32 | 319 | 0.38 | 0.35 | 108 | 90.1 | 0:10:09 | 8,166 | 9.74 |
| *H. sapiens* chr. 18 | 12× | 922 | 5.37 | 277 | 1.26 | 1.25 | 86 | 94.3 | 0:27:47 | 24,309 | 14.52 |
|  | Top | 2,282 | 2.86 | 452 | 0.84 | 0.72 | 119 | 93.8 | 1:02:50 | 54,787 | 23.75 |
|  | Both | 2,588 | 2.13 | 424 | 0.79 | 0.77 | 118 | 95.7 | 3:21:02 | 174,815 | 47.26 |

**Table S8:**
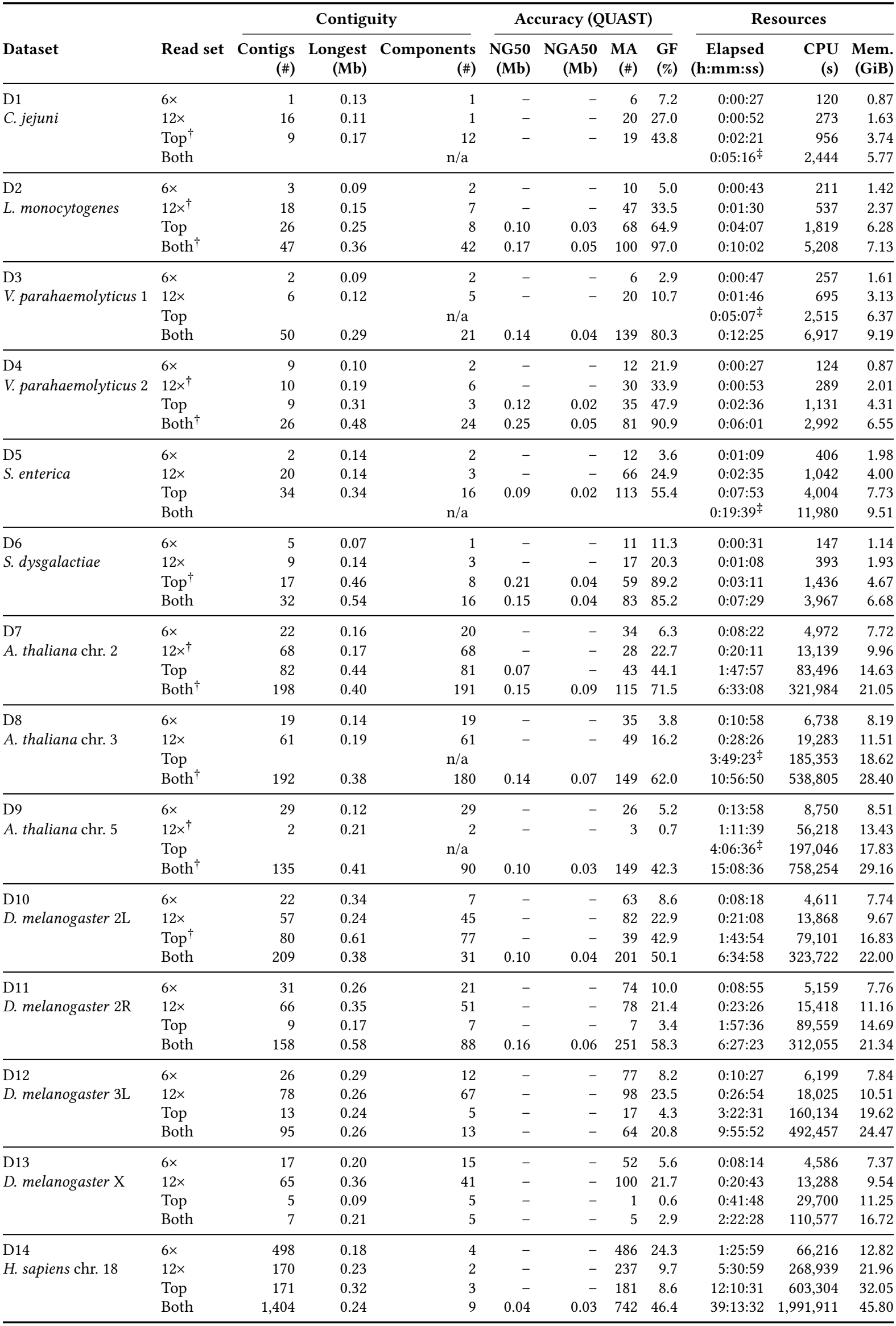
Assemblies by Rawsamble. † Rawsamble retried by limiting overlaps per read. n/a, ‡ miniasm cannot generate the contig sequences of Rawsamble.

|  |  | Contiguity |  |  | Accuracy (QUAST) |  |  |  | Resources |  |  |
|---|---|---|---|---|---|---|---|---|---|---|---|
| Dataset | Read set | Contigs<br>(#) | Longest<br>(Mb) | Components<br>(#) | NG50<br>(Mb) | NGA50<br>(Mb) | MA<br>(#) | GF<br>(%) | Elapsed<br>(h:mm:ss) | CPU<br>(s) | Mem.<br>(GiB) |
| D1 | 6× | 1 | 0.13 | 1 | – | – | 6 | 7.2 | 0:00:27 | 120 | 0.87 |
| *C. jejuni* | 12× | 16 | 0.11 | 1 | – | – | 20 | 27.0 | 0:00:52 | 273 | 1.63 |
|  | Top<sup>†</sup> | 9 | 0.17 | 12 | – | – | 19 | 43.8 | 0:02:21 | 956 | 3.74 |
|  | Both | n/a |  |  |  |  |  |  | 0:05:16<sup>‡</sup> | 2,444 | 5.77 |
| D2 | 6× | 3 | 0.09 | 2 | – | – | 10 | 5.0 | 0:00:43 | 211 | 1.42 |
| *L. monocytogenes* | 12×<sup>†</sup> | 18 | 0.15 | 7 | – | – | 47 | 33.5 | 0:01:30 | 537 | 2.37 |
|  | Top | 26 | 0.25 | 8 | 0.10 | 0.03 | 68 | 64.9 | 0:04:07 | 1,819 | 6.28 |
|  | Both<sup>†</sup> | 47 | 0.36 | 42 | 0.17 | 0.05 | 100 | 97.0 | 0:10:02 | 5,208 | 7.13 |
| D3 | 6× | 2 | 0.09 | 2 | – | – | 6 | 2.9 | 0:00:47 | 257 | 1.61 |
| *V. parahaemolyticus* 1 | 12× | 6 | 0.12 | 5 | – | – | 20 | 10.7 | 0:01:46 | 695 | 3.13 |
|  | Top | n/a |  |  |  |  |  |  | 0:05:07<sup>‡</sup> | 2,515 | 6.37 |
|  | Both | 50 | 0.29 | 21 | 0.14 | 0.04 | 139 | 80.3 | 0:12:25 | 6,917 | 9.19 |
| D4 | 6× | 9 | 0.10 | 2 | – | – | 12 | 21.9 | 0:00:27 | 124 | 0.87 |
| *V. parahaemolyticus* 2 | 12×<sup>†</sup> | 10 | 0.19 | 6 | – | – | 30 | 33.9 | 0:00:53 | 289 | 2.01 |
|  | Top | 9 | 0.31 | 3 | 0.12 | 0.02 | 35 | 47.9 | 0:02:36 | 1,131 | 4.31 |
|  | Both<sup>†</sup> | 26 | 0.48 | 24 | 0.25 | 0.05 | 81 | 90.9 | 0:06:01 | 2,992 | 6.55 |
| D5 | 6× | 2 | 0.14 | 2 | – | – | 12 | 3.6 | 0:01:09 | 406 | 1.98 |
| *S. enterica* | 12× | 20 | 0.14 | 3 | – | – | 66 | 24.9 | 0:02:35 | 1,042 | 4.00 |
|  | Top | 34 | 0.34 | 16 | 0.09 | 0.02 | 113 | 55.4 | 0:07:53 | 4,004 | 7.73 |
|  | Both | n/a |  |  |  |  |  |  | 0:19:39<sup>‡</sup> | 11,980 | 9.51 |
| D6 | 6× | 5 | 0.07 | 1 | – | – | 11 | 11.3 | 0:00:31 | 147 | 1.14 |
| *S. dysgalactiae* | 12× | 9 | 0.14 | 3 | – | – | 17 | 20.3 | 0:01:08 | 393 | 1.93 |
|  | Top<sup>†</sup> | 17 | 0.46 | 8 | 0.21 | 0.04 | 59 | 89.2 | 0:03:11 | 1,436 | 4.67 |
|  | Both | 32 | 0.54 | 16 | 0.15 | 0.04 | 83 | 85.2 | 0:07:29 | 3,967 | 6.68 |
| D7 | 6× | 22 | 0.16 | 20 | – | – | 34 | 6.3 | 0:08:22 | 4,972 | 7.72 |
| *A. thaliana* chr. 2 | 12×<sup>†</sup> | 68 | 0.17 | 68 | – | – | 28 | 22.7 | 0:20:11 | 13,139 | 9.96 |
|  | Top | 82 | 0.44 | 81 | 0.07 | – | 43 | 44.1 | 1:47:57 | 83,496 | 14.63 |
|  | Both<sup>†</sup> | 198 | 0.40 | 191 | 0.15 | 0.09 | 115 | 71.5 | 6:33:08 | 321,984 | 21.05 |
| D8 | 6× | 19 | 0.14 | 19 | – | – | 35 | 3.8 | 0:10:58 | 6,738 | 8.19 |
| *A. thaliana* chr. 3 | 12× | 61 | 0.19 | 61 | – | – | 49 | 16.2 | 0:28:26 | 19,283 | 11.51 |
|  | Top | n/a |  |  |  |  |  |  | 3:49:23<sup>‡</sup> | 185,353 | 18.62 |
|  | Both<sup>†</sup> | 192 | 0.38 | 180 | 0.14 | 0.07 | 149 | 62.0 | 10:56:50 | 538,805 | 28.40 |
| D9 | 6× | 29 | 0.12 | 29 | – | – | 26 | 5.2 | 0:13:58 | 8,750 | 8.51 |
| *A. thaliana* chr. 5 | 12×<sup>†</sup> | 2 | 0.21 | 2 | – | – | 3 | 0.7 | 1:11:39 | 56,218 | 13.43 |
|  | Top | n/a |  |  |  |  |  |  | 4:06:36<sup>‡</sup> | 197,046 | 17.83 |
|  | Both<sup>†</sup> | 135 | 0.41 | 90 | 0.10 | 0.03 | 149 | 42.3 | 15:08:36 | 758,254 | 29.16 |
| D10 | 6× | 22 | 0.34 | 7 | – | – | 63 | 8.6 | 0:08:18 | 4,611 | 7.74 |
| *D. melanogaster* 2L | 12× | 57 | 0.24 | 45 | – | – | 82 | 22.9 | 0:21:08 | 13,868 | 9.67 |
|  | Top<sup>†</sup> | 80 | 0.61 | 77 | – | – | 39 | 42.9 | 1:43:54 | 79,101 | 16.83 |
|  | Both | 209 | 0.38 | 31 | 0.10 | 0.04 | 201 | 50.1 | 6:34:58 | 323,722 | 22.00 |
| D11 | 6× | 31 | 0.26 | 21 | – | – | 74 | 10.0 | 0:08:55 | 5,159 | 7.76 |
| *D. melanogaster* 2R | 12× | 66 | 0.35 | 51 | – | – | 78 | 21.4 | 0:23:26 | 15,418 | 11.16 |
|  | Top | 9 | 0.17 | 7 | – | – | 7 | 3.4 | 1:57:36 | 89,559 | 14.69 |
|  | Both | 158 | 0.58 | 88 | 0.16 | 0.06 | 251 | 58.3 | 6:27:23 | 312,055 | 21.34 |
| D12 | 6× | 26 | 0.29 | 12 | – | – | 77 | 8.2 | 0:10:27 | 6,199 | 7.84 |
| *D. melanogaster* 3L | 12× | 78 | 0.26 | 67 | – | – | 98 | 23.5 | 0:26:54 | 18,025 | 10.51 |
|  | Top | 13 | 0.24 | 5 | – | – | 17 | 4.3 | 3:22:31 | 160,134 | 19.62 |
|  | Both | 95 | 0.26 | 13 | – | – | 64 | 20.8 | 9:55:52 | 492,457 | 24.47 |
| D13 | 6× | 17 | 0.20 | 15 | – | – | 52 | 5.6 | 0:08:14 | 4,586 | 7.37 |
| *D. melanogaster* X | 12× | 65 | 0.36 | 41 | – | – | 100 | 21.7 | 0:20:43 | 13,288 | 9.54 |
|  | Top | 5 | 0.09 | 5 | – | – | 1 | 0.6 | 0:41:48 | 29,700 | 11.25 |
|  | Both | 7 | 0.21 | 5 | – | – | 5 | 2.9 | 2:22:28 | 110,577 | 16.72 |
| D14 | 6× | 498 | 0.18 | 4 | – | – | 486 | 24.3 | 1:25:59 | 66,216 | 12.82 |
| *H. sapiens* chr. 18 | 12× | 170 | 0.23 | 2 | – | – | 237 | 9.7 | 5:30:59 | 268,939 | 21.96 |
|  | Top | 171 | 0.32 | 3 | – | – | 181 | 8.6 | 12:10:31 | 603,304 | 32.05 |
|  | Both | 1,404 | 0.24 | 9 | 0.04 | 0.03 | 742 | 46.4 | 39:13:32 | 1,991,911 | 45.80 |

**Table S9:** Assemblies by Flye.

|  |  | **Contiguity** |  |  | **Accuracy (QUAST)** |  |  |  | **Resources** |  |  |
|---|---|---|---|---|---|---|---|---|---|---|---|
| **Dataset** | **Read set** | **Contigs<br>(#)** | **Longest<br>(Mb)** | **Components<br>(#)** | **NG50<br>(Mb)** | **NGA50<br>(Mb)** | **MA<br>(#)** | **GF<br>(%)** | **Elapsed<br>(h:mm:ss)** | **CPU<br>(s)** | **Mem.<br>(GiB)** |
| D1<br>*C. jejuni* | 6× | 8 | 0.81 | 13 | 0.30 | 0.30 | 0 | 98.7 | 0:01:12 | 216 | 0.59 |
|  | 12× | 1 | 1.77 | 2 | 1.77 | 1.77 | 0 | 100.0 | 0:01:55 | 256 | 0.72 |
|  | Top | 1 | 1.77 | 1 | 1.77 | 1.77 | 0 | 100.0 | 0:02:40 | 453 | 1.31 |
|  | Both | 1 | 1.77 | 1 | 1.77 | 1.77 | 0 | 100.0 | 0:04:42 | 832 | 1.64 |
| D2<br>*L. monocytogenes* | 6× | 22 | 0.42 | 23 | 0.21 | 0.21 | 2 | 97.1 | 0:00:49 | 268 | 0.51 |
|  | 12× | 1 | 2.94 | 1 | 2.94 | 2.94 | 0 | 100.0 | 0:03:58 | 457 | 0.85 |
|  | Top | 1 | 2.94 | 1 | 2.94 | 2.94 | 0 | 100.0 | 0:03:53 | 698 | 1.48 |
|  | Both | 1 | 2.94 | 1 | 2.94 | 2.94 | 0 | 100.0 | 0:14:46 | 1,831 | 2.34 |
| D3<br>*V. parahaemolyticus* 1 | 6× | 17 | 0.46 | 15 | 0.22 | 0.22 | 2 | 97.4 | 0:03:41 | 466 | 0.49 |
|  | 12× | 2 | 3.16 | 1 | 3.16 | 3.16 | 0 | 99.9 | 0:17:03 | 2,230 | 0.72 |
|  | Top | 1 | 3.28 | 2 | 3.28 | 3.28 | 0 | 100.0 | 0:13:59 | 1,667 | 1.45 |
|  | Both | 2 | 3.28 | 2 | 3.28 | 3.28 | 0 | 100.0 | 0:08:22 | 2,406 | 1.81 |
| D4<br>*V. parahaemolyticus* 2 | 6× | 9 | 1.12 | 9 | 1.12 | 1.12 | 1 | 90.2 | 0:00:49 | 150 | 0.45 |
|  | 12× | 1 | 1.87 | 1 | 1.87 | 1.87 | 0 | 100.0 | 0:01:19 | 216 | 0.75 |
|  | Top | 1 | 1.87 | 1 | 1.87 | 1.87 | 0 | 100.0 | 0:02:24 | 418 | 0.91 |
|  | Both | 1 | 1.87 | 1 | 1.87 | 1.87 | 0 | 100.0 | 0:04:15 | 770 | 1.29 |
| D5<br>*S. enterica* | 6× | 35 | 0.49 | 31 | 0.18 | 0.15 | 8 | 96.3 | 0:00:58 | 340 | 0.49 |
|  | 12× | 3 | 2.74 | 6 | 2.74 | 2.73 | 1 | 100.0 | 0:02:26 | 543 | 0.59 |
|  | Top | 1 | 4.75 | 1 | 4.75 | 4.75 | 0 | 100.0 | 0:09:43 | 1,915 | 1.19 |
|  | Both | 1 | 4.75 | 1 | 4.75 | 4.75 | 0 | 100.0 | 0:15:09 | 3,407 | 1.91 |
| D6<br>*S. dysgalactiae* | 6× | 12 | 0.71 | 12 | 0.28 | 0.28 | 1 | 97.9 | 0:02:23 | 306 | 0.48 |
|  | 12× | 1 | 2.21 | 1 | 2.21 | 2.21 | 0 | 100.0 | 0:01:20 | 265 | 0.77 |
|  | Top | 1 | 2.21 | 1 | 2.21 | 2.21 | 0 | 100.0 | 0:02:18 | 554 | 0.97 |
|  | Both | 1 | 2.20 | 1 | 2.20 | 2.20 | 0 | 100.0 | 0:05:28 | 1,121 | 1.36 |
| D7<br>*A. thaliana* chr. 2 | 6× | 31 | 3.73 | 26 | 0.92 | 0.92 | 12 | 98.8 | 0:05:07 | 1,733 | 2.77 |
|  | 12× | 7 | 16.31 | 8 | 16.31 | 16.11 | 13 | 100.0 | 0:08:12 | 3,687 | 2.97 |
|  | Top | 11 | 16.22 | 6 | 16.22 | 16.12 | 8 | 100.0 | 0:14:16 | 7,628 | 5.01 |
|  | Both | 8 | 14.51 | 3 | 14.51 | 14.50 | 14 | 100.0 | 0:24:50 | 14,032 | 6.93 |
| D8<br>*A. thaliana* chr. 3 | 6× | 50 | 2.33 | 50 | 0.75 | 0.75 | 11 | 96.8 | 0:08:30 | 4,617 | 7.62 |
|  | 12× | 13 | 9.34 | 10 | 7.11 | 7.11 | 17 | 99.6 | 0:18:44 | 11,857 | 11.59 |
|  | Top | 10 | 13.70 | 7 | 13.70 | 13.59 | 16 | 99.6 | 0:51:11 | 38,652 | 14.02 |
|  | Both | 8 | 13.70 | 5 | 13.70 | 13.59 | 18 | 99.6 | 6:45:48 | 101,170 | 14.36 |
| D9<br>*A. thaliana* chr. 5 | 6× | 45 | 2.69 | 42 | 1.23 | 1.23 | 12 | 98.9 | 0:06:44 | 2,966 | 3.89 |
|  | 12× | 29 | 10.38 | 29 | 8.90 | 8.84 | 12 | 99.7 | 8:06:17 | 33,802 | 5.46 |
|  | Top | 19 | 11.24 | 21 | 8.97 | 7.73 | 12 | 99.7 | 0:21:09 | 13,103 | 7.82 |
|  | Both | 20 | 11.25 | 17 | 6.36 | 6.35 | 15 | 99.7 | 0:34:11 | 22,442 | 9.69 |
| D10<br>*D. melanogaster* 2L | 6× | 87 | 1.36 | 88 | 0.40 | 0.40 | 8 | 91.1 | 0:03:49 | 1,837 | 4.52 |
|  | 12× | 13 | 5.69 | 13 | 3.44 | 3.39 | 8 | 98.9 | 0:07:08 | 3,187 | 6.65 |
|  | Top | 4 | 21.48 | 2 | 21.48 | 15.73 | 4 | 99.4 | 0:11:34 | 6,082 | 8.79 |
|  | Both | 3 | 21.55 | 2 | 21.55 | 15.76 | 4 | 99.6 | 0:19:08 | 10,402 | 10.37 |
| D11<br>*D. melanogaster* 2R | 6× | 89 | 1.26 | 94 | 0.50 | 0.45 | 22 | 85.8 | 0:04:02 | 2,159 | 5.34 |
|  | 12× | 38 | 9.40 | 32 | 3.61 | 3.60 | 34 | 96.8 | 0:06:53 | 3,252 | 6.05 |
|  | Top | 14 | 21.41 | 20 | 21.41 | 21.39 | 23 | 98.9 | 0:21:27 | 13,382 | 9.39 |
|  | Both | 17 | 21.65 | 21 | 21.65 | 21.62 | 15 | 99.1 | 0:28:36 | 18,323 | 10.21 |
| D12<br>*D. melanogaster* 3L | 6× | 96 | 1.22 | 93 | 0.38 | 0.36 | 28 | 87.2 | 0:05:28 | 2,861 | 6.43 |
|  | 12× | 46 | 4.48 | 34 | 1.58 | 1.58 | 12 | 97.8 | 0:06:47 | 3,623 | 7.25 |
|  | Top | 17 | 16.98 | 14 | 16.98 | 16.98 | 17 | 99.2 | 0:13:10 | 7,172 | 9.87 |
|  | Both | 12 | 24.44 | 15 | 24.44 | 24.44 | 23 | 99.3 | 0:23:44 | 14,064 | 11.71 |
| D13<br>*D. melanogaster* X | 6× | 116 | 0.72 | 117 | 0.35 | 0.35 | 45 | 92.2 | 0:04:44 | 2,528 | 6.79 |
|  | 12× | 50 | 4.42 | 39 | 1.73 | 1.73 | 52 | 98.0 | 0:10:05 | 4,298 | 9.37 |
|  | Top | 43 | 7.62 | 30 | 6.44 | 6.44 | 87 | 99.1 | 0:11:24 | 5,895 | 10.42 |
|  | Both | 31 | 16.64 | 24 | 16.64 | 16.64 | 99 | 99.4 | 0:18:16 | 9,915 | 11.07 |
| D14<br>*H. sapiens* chr. 18 | 6× | 190 | 1.71 | 228 | 0.63 | 0.61 | 84 | 92.2 | 0:12:01 | 7,121 | 8.76 |
|  | 12× | 40 | 13.75 | 48 | 7.44 | 7.41 | 16 | 95.3 | 0:19:40 | 12,346 | 7.38 |
|  | Top | 28 | 33.14 | 19 | 11.66 | 11.65 | 8 | 95.4 | 0:30:14 | 20,123 | 6.48 |
|  | Both | 41 | 33.14 | 19 | 26.04 | 26.04 | 13 | 95.5 | 1:03:27 | 45,351 | 7.27 |

**Table S10:**
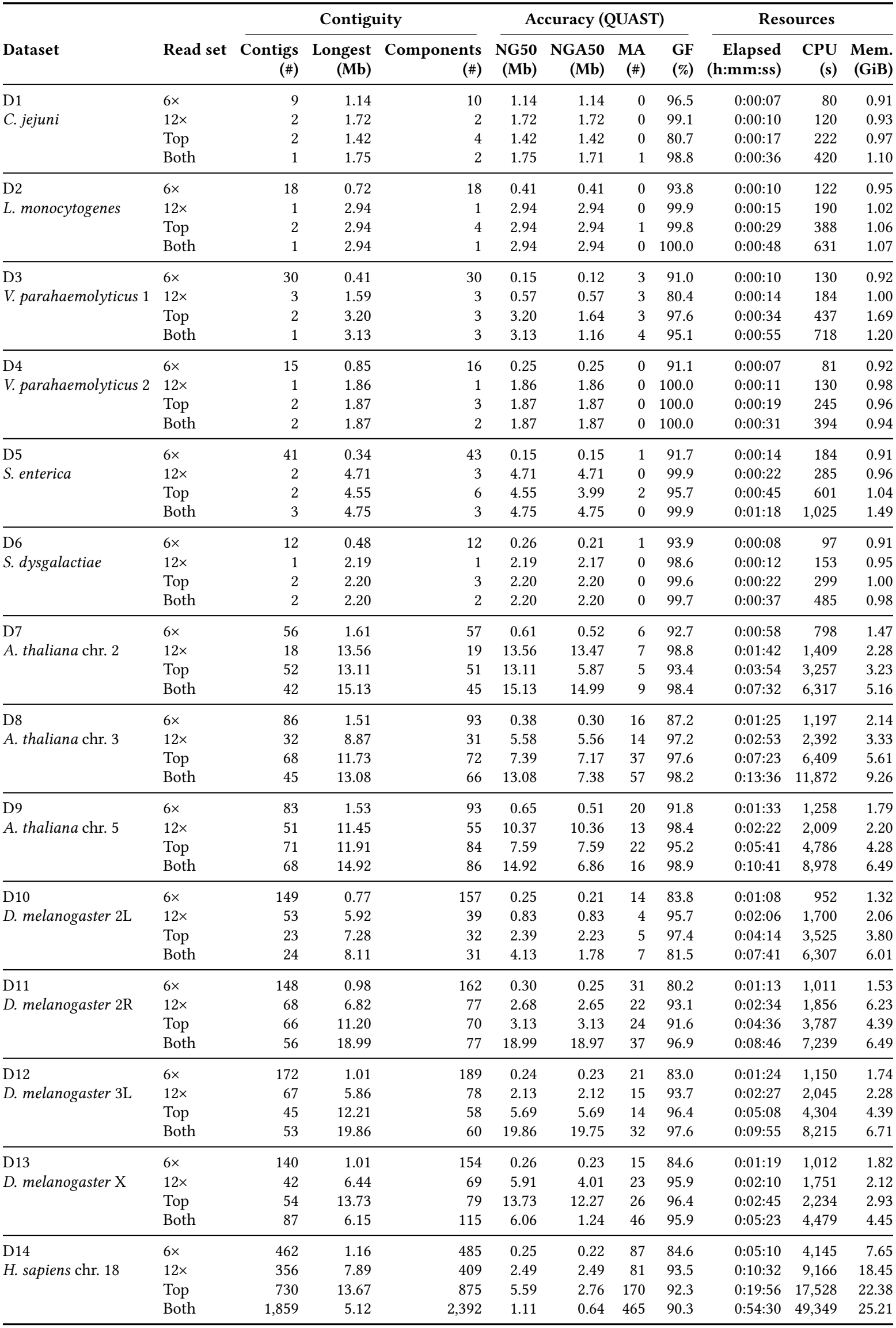
Assemblies by wtdbg2.

| Dataset | Read set | Contiguity |  |  | Accuracy (QUAST) |  |  |  | Resources |  |  |
|---|---|---|---|---|---|---|---|---|---|---|---|
|  |  | **Contigs**<br>**(#)** | **Longest**<br>**(Mb)** | **Components**<br>**(#)** | **NG50**<br>**(Mb)** | **NGA50**<br>**(Mb)** | **MA**<br>**(#)** | **GF**<br>**(%)** | **Elapsed**<br>**(h:mm:ss)** | **CPU**<br>**(s)** | **Mem.**<br>**(GiB)** |
| D1<br>*C. jejuni* | 6× | 9 | 1.14 | 10 | 1.14 | 1.14 | 0 | 96.5 | 0:00:07 | 80 | 0.91 |
|  | 12× | 2 | 1.72 | 2 | 1.72 | 1.72 | 0 | 99.1 | 0:00:10 | 120 | 0.93 |
|  | Top | 2 | 1.42 | 4 | 1.42 | 1.42 | 0 | 80.7 | 0:00:17 | 222 | 0.97 |
|  | Both | 1 | 1.75 | 2 | 1.75 | 1.71 | 1 | 98.8 | 0:00:36 | 420 | 1.10 |
| D2<br>*L. monocytogenes* | 6× | 18 | 0.72 | 18 | 0.41 | 0.41 | 0 | 93.8 | 0:00:10 | 122 | 0.95 |
|  | 12× | 1 | 2.94 | 1 | 2.94 | 2.94 | 0 | 99.9 | 0:00:15 | 190 | 1.02 |
|  | Top | 2 | 2.94 | 4 | 2.94 | 2.94 | 1 | 99.8 | 0:00:29 | 388 | 1.06 |
|  | Both | 1 | 2.94 | 1 | 2.94 | 2.94 | 0 | 100.0 | 0:00:48 | 631 | 1.07 |
| D3<br>*V. parahaemolyticus* 1 | 6× | 30 | 0.41 | 30 | 0.15 | 0.12 | 3 | 91.0 | 0:00:10 | 130 | 0.92 |
|  | 12× | 3 | 1.59 | 3 | 0.57 | 0.57 | 3 | 80.4 | 0:00:14 | 184 | 1.00 |
|  | Top | 2 | 3.20 | 3 | 3.20 | 1.64 | 3 | 97.6 | 0:00:34 | 437 | 1.69 |
|  | Both | 1 | 3.13 | 3 | 3.13 | 1.16 | 4 | 95.1 | 0:00:55 | 718 | 1.20 |
| D4<br>*V. parahaemolyticus* 2 | 6× | 15 | 0.85 | 16 | 0.25 | 0.25 | 0 | 91.1 | 0:00:07 | 81 | 0.92 |
|  | 12× | 1 | 1.86 | 1 | 1.86 | 1.86 | 0 | 100.0 | 0:00:11 | 130 | 0.98 |
|  | Top | 2 | 1.87 | 3 | 1.87 | 1.87 | 0 | 100.0 | 0:00:19 | 245 | 0.96 |
|  | Both | 2 | 1.87 | 2 | 1.87 | 1.87 | 0 | 100.0 | 0:00:31 | 394 | 0.94 |
| D5<br>*S. enterica* | 6× | 41 | 0.34 | 43 | 0.15 | 0.15 | 1 | 91.7 | 0:00:14 | 184 | 0.91 |
|  | 12× | 2 | 4.71 | 3 | 4.71 | 4.71 | 0 | 99.9 | 0:00:22 | 285 | 0.96 |
|  | Top | 2 | 4.55 | 6 | 4.55 | 3.99 | 2 | 95.7 | 0:00:45 | 601 | 1.04 |
|  | Both | 3 | 4.75 | 3 | 4.75 | 4.75 | 0 | 99.9 | 0:01:18 | 1,025 | 1.49 |
| D6<br>*S. dysgalactiae* | 6× | 12 | 0.48 | 12 | 0.26 | 0.21 | 1 | 93.9 | 0:00:08 | 97 | 0.91 |
|  | 12× | 1 | 2.19 | 1 | 2.19 | 2.17 | 0 | 98.6 | 0:00:12 | 153 | 0.95 |
|  | Top | 2 | 2.20 | 3 | 2.20 | 2.20 | 0 | 99.6 | 0:00:22 | 299 | 1.00 |
|  | Both | 2 | 2.20 | 2 | 2.20 | 2.20 | 0 | 99.7 | 0:00:37 | 485 | 0.98 |
| D7<br>*A. thaliana* chr. 2 | 6× | 56 | 1.61 | 57 | 0.61 | 0.52 | 6 | 92.7 | 0:00:58 | 798 | 1.47 |
|  | 12× | 18 | 13.56 | 19 | 13.56 | 13.47 | 7 | 98.8 | 0:01:42 | 1,409 | 2.28 |
|  | Top | 52 | 13.11 | 51 | 13.11 | 5.87 | 5 | 93.4 | 0:03:54 | 3,257 | 3.23 |
|  | Both | 42 | 15.13 | 45 | 15.13 | 14.99 | 9 | 98.4 | 0:07:32 | 6,317 | 5.16 |
| D8<br>*A. thaliana* chr. 3 | 6× | 86 | 1.51 | 93 | 0.38 | 0.30 | 16 | 87.2 | 0:01:25 | 1,197 | 2.14 |
|  | 12× | 32 | 8.87 | 31 | 5.58 | 5.56 | 14 | 97.2 | 0:02:53 | 2,392 | 3.33 |
|  | Top | 68 | 11.73 | 72 | 7.39 | 7.17 | 37 | 97.6 | 0:07:23 | 6,409 | 5.61 |
|  | Both | 45 | 13.08 | 66 | 13.08 | 7.38 | 57 | 98.2 | 0:13:36 | 11,872 | 9.26 |
| D9<br>*A. thaliana* chr. 5 | 6× | 83 | 1.53 | 93 | 0.65 | 0.51 | 20 | 91.8 | 0:01:33 | 1,258 | 1.79 |
|  | 12× | 51 | 11.45 | 55 | 10.37 | 10.36 | 13 | 98.4 | 0:02:22 | 2,009 | 2.20 |
|  | Top | 71 | 11.91 | 84 | 7.59 | 7.59 | 22 | 95.2 | 0:05:41 | 4,786 | 4.28 |
|  | Both | 68 | 14.92 | 86 | 14.92 | 6.86 | 16 | 98.9 | 0:10:41 | 8,978 | 6.49 |
| D10<br>*D. melanogaster* 2L | 6× | 149 | 0.77 | 157 | 0.25 | 0.21 | 14 | 83.8 | 0:01:08 | 952 | 1.32 |
|  | 12× | 53 | 5.92 | 39 | 0.83 | 0.83 | 4 | 95.7 | 0:02:06 | 1,700 | 2.06 |
|  | Top | 23 | 7.28 | 32 | 2.39 | 2.23 | 5 | 97.4 | 0:04:14 | 3,525 | 3.80 |
|  | Both | 24 | 8.11 | 31 | 4.13 | 1.78 | 7 | 81.5 | 0:07:41 | 6,307 | 6.01 |
| D11<br>*D. melanogaster* 2R | 6× | 148 | 0.98 | 162 | 0.30 | 0.25 | 31 | 80.2 | 0:01:13 | 1,011 | 1.53 |
|  | 12× | 68 | 6.82 | 77 | 2.68 | 2.65 | 22 | 93.1 | 0:02:34 | 1,856 | 6.23 |
|  | Top | 66 | 11.20 | 70 | 3.13 | 3.13 | 24 | 91.6 | 0:04:36 | 3,787 | 4.39 |
|  | Both | 56 | 18.99 | 77 | 18.99 | 18.97 | 37 | 96.9 | 0:08:46 | 7,239 | 6.49 |
| D12<br>*D. melanogaster* 3L | 6× | 172 | 1.01 | 189 | 0.24 | 0.23 | 21 | 83.0 | 0:01:24 | 1,150 | 1.74 |
|  | 12× | 67 | 5.86 | 78 | 2.13 | 2.12 | 15 | 93.7 | 0:02:27 | 2,045 | 2.28 |
|  | Top | 45 | 12.21 | 58 | 5.69 | 5.69 | 14 | 96.4 | 0:05:08 | 4,304 | 4.39 |
|  | Both | 53 | 19.86 | 60 | 19.86 | 19.75 | 32 | 97.6 | 0:09:55 | 8,215 | 6.71 |
| D13<br>*D. melanogaster* X | 6× | 140 | 1.01 | 154 | 0.26 | 0.23 | 15 | 84.6 | 0:01:19 | 1,012 | 1.82 |
|  | 12× | 42 | 6.44 | 69 | 5.91 | 4.01 | 23 | 95.9 | 0:02:10 | 1,751 | 2.12 |
|  | Top | 54 | 13.73 | 79 | 13.73 | 12.27 | 26 | 96.4 | 0:02:45 | 2,234 | 2.93 |
|  | Both | 87 | 6.15 | 115 | 6.06 | 1.24 | 46 | 95.9 | 0:05:23 | 4,479 | 4.45 |
| D14<br>*H. sapiens* chr. 18 | 6× | 462 | 1.16 | 485 | 0.25 | 0.22 | 87 | 84.6 | 0:05:10 | 4,145 | 7.65 |
|  | 12× | 356 | 7.89 | 409 | 2.49 | 2.49 | 81 | 93.5 | 0:10:32 | 9,166 | 18.45 |
|  | Top | 730 | 13.67 | 875 | 5.59 | 2.76 | 170 | 92.3 | 0:19:56 | 17,528 | 22.38 |
|  | Both | 1,859 | 5.12 | 2,392 | 1.11 | 0.64 | 465 | 90.3 | 0:54:30 | 49,349 | 25.21 |

#### S4.1. Bandage plots

In Supplementary Figures S1 and S2 we show the Bandage [13] plots of the assemblies generated by all tools on the top-depth forward strand datasets.

#### S4.2. Nucleotide assignment verification

HARICOT as a raw signal tool does not support outputting nucleotide sequences, which are however required for QUAST [11] analysis. To assign nucleotide sequences to nodes in the assembly graph, we map the coordinates of the anchor read from event space to nucleotide space using the move table generated during basecalling by Dorado [5]. Each node is then assigned the nucleotide sequence of its anchor read up to the next node. To verify this approach, we apply the same method to the assembly graphs of wtdbg2 [4] and compare with wtdbg2’s consensus using QUAST. Supplementary Table S11 shows the results. As expected, NGA50 suffers, since QUAST breaks alignments at misassemblies. In most cases, the other metrics provide a reasonable approximation of the consensus.

**Table S11:**
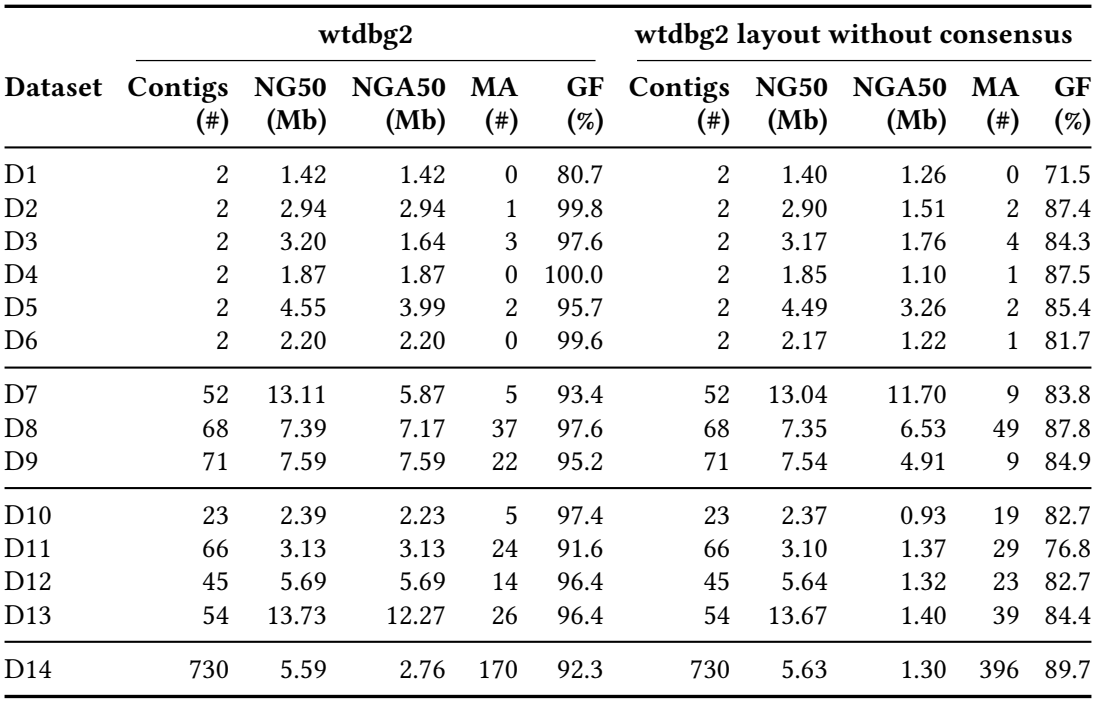
wtdbg2 with and without consensus, forward-strand datasets at the top depth.

**Figure S1:**
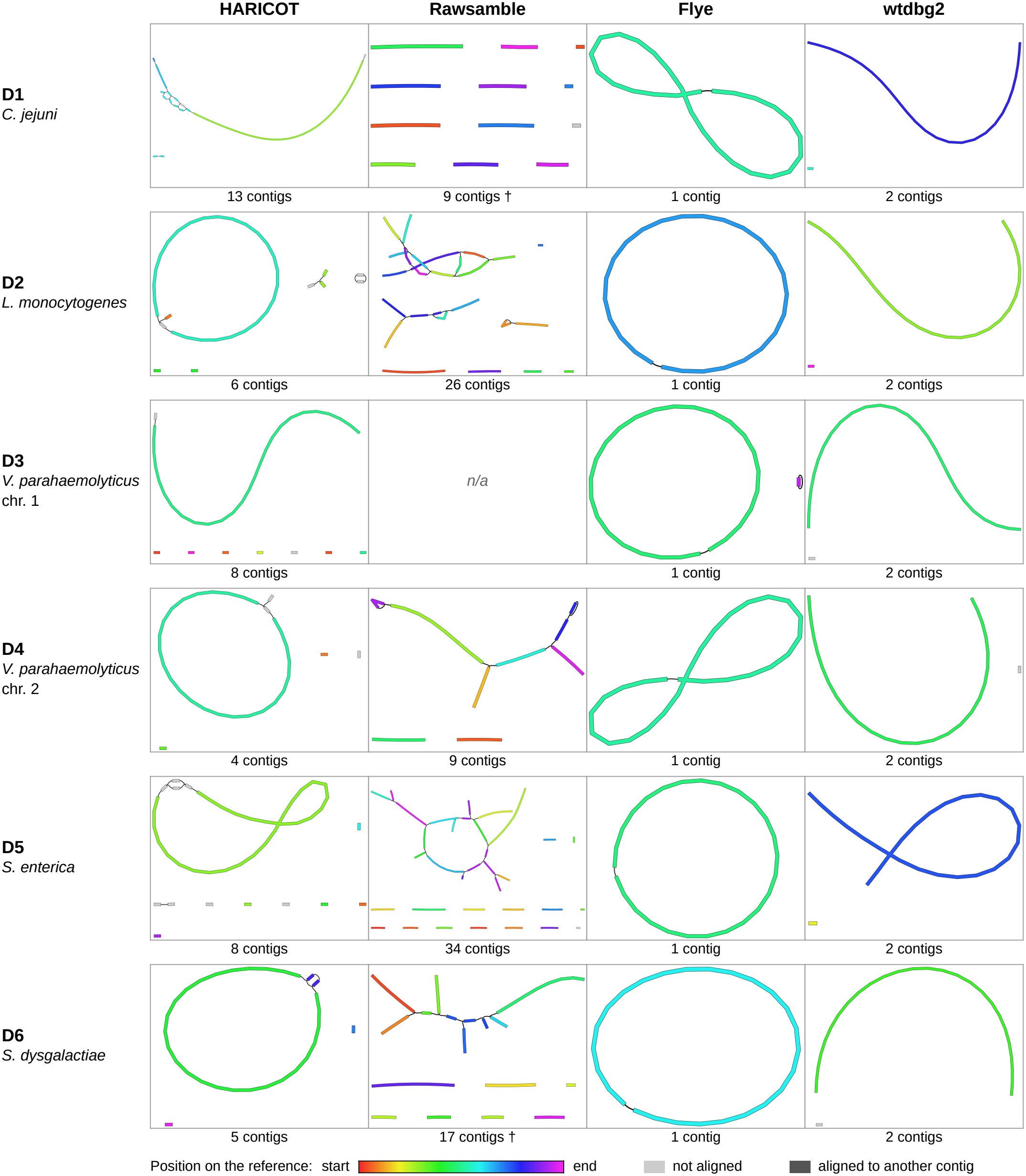
Bandage plots on top-depth bacterial forward strand datasets. † Rawsamble retried by limiting overlaps per read.

**Figure S2:**
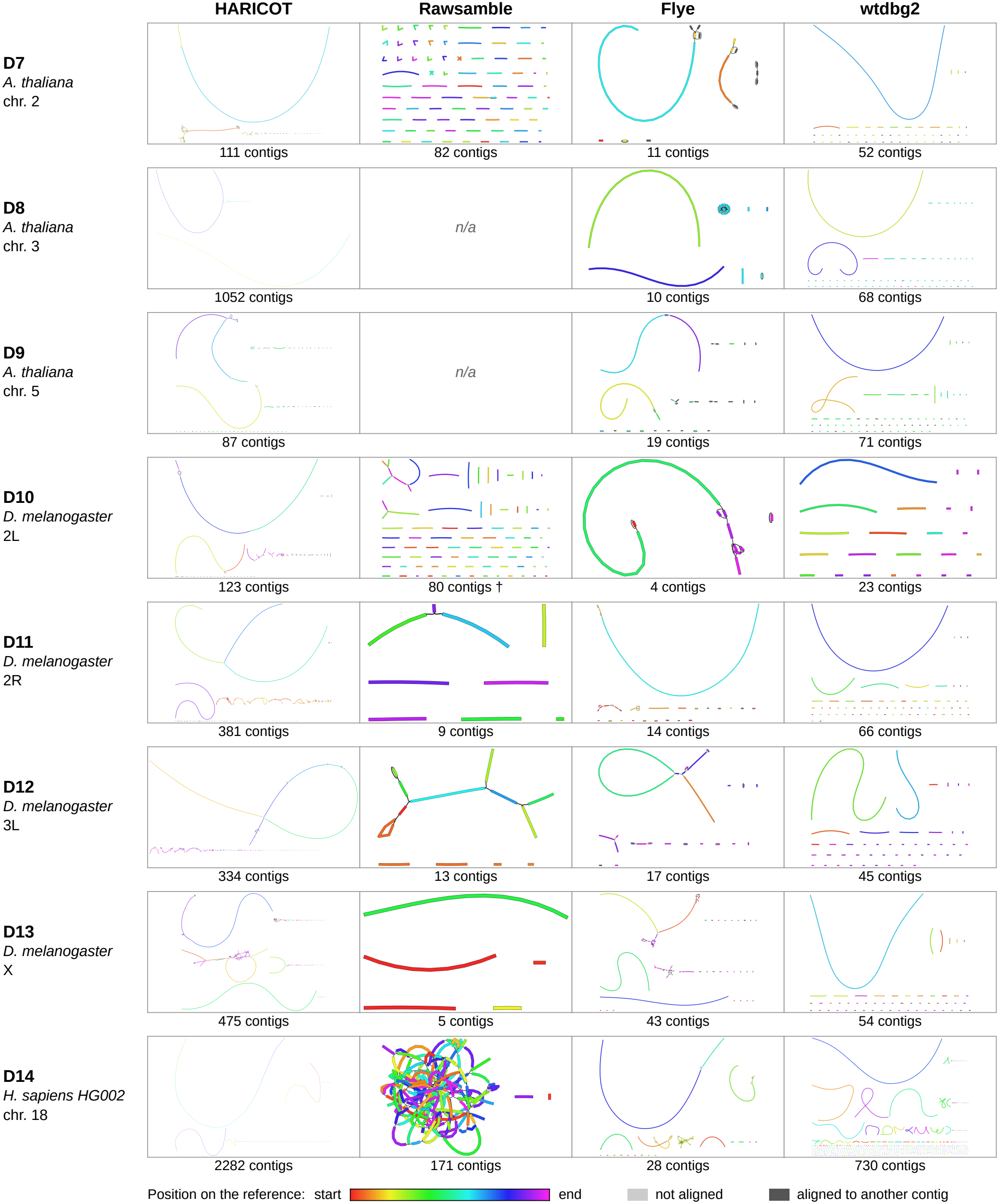
Bandage plots on top-depth eukaryotic forward strand datasets. † Rawsamble retried by limiting overlaps per read.

### S5. Algorithms

#### S5.1. Overlap classifier features

Supplementary Table S12 lists the features of the three overlap classifiers, stage 1 (S1), stage 2 (S2) and the contig-end rescue (R), each an ensemble of five 400-tree random forests. *A* is the longer read of a pair.

**Table S12:** Features of the overlap classifiers.

| Feature | Description | S1 | S2 | R |
|---|---|---|---|---|
| z | significance of the seed diagonal | • | • |  |
| slope, abs_log_slope | path slope; \|ln slope\| | • |  |  |
| a_bins, b_bins | read lengths | • | • | • |
| span_a, ov_frac_short | overlap length on *A*; as a fraction of the shorter read | • | • | • |
| oh_left, oh_right, oh_frac | overhang shared by both reads | • | • | • |
| a_contained, b_contained | read contained in the overlap | • |  | • |
| hole_frac | share of path bins without seeds | • | • |  |
| max_hole | longest gap in the path | • |  |  |
| resid_med, _max, _frac1 | path residuals from the fitted line | • |  |  |
| anc_per_bin | seeds on the fitted line per bin | • | • | • |
| dens_cv, bins_hit | unevenness and coverage of those seeds | • |  | • |
| high_occ_frac, occ_rel_med | share of frequent seeds; median seed occurrence | • |  | • |
| occ_rel_q90 | 90th percentile of seed occurrence | • | • | • |
| ctx_rel_a, ctx_rel_b | candidate depth over the overlap, relative to the read | • |  | • |
| ctx_abs | the same, relative to the dataset | • | • | • |
| ctx_pois | Poisson tail of that depth at the read’s depth | • | • | • |
| ctx_pois_abs | the same, at the dataset’s depth | • |  | • |
| deg_rel | candidate partners per bin, relative to the dataset | • | • | • |
| tri_frac, tri_rel | common candidate partners with consistent offsets | • | • | • |
| dtw_mean, dtw_decoy | signal DTW cost along the overlap; at a decoy offset | • |  | • |
| dtw_ratio | dtw_mean / dtw_decoy | • | • | • |
| dtw_good_frac | share of windows with a low DTW ratio | • |  | • |
| dtw_r_q10, _q50, _q90, _max | quantiles of the per-window DTW ratio | • |  | • |
| s1 | stage-1 score |  | • |  |
| mut_a, mut_b | accepted partners shared by *A* and *B* |  | • |  |
| cons_a, cons_b | of those, with consistent offsets |  | • |  |
| nbr_a, nbr_b | accepted partners, relative to the dataset |  | • |  |
| nb_score | mean stage-1 score of the partners |  | • |  |
| rank_a, rank_b | rank of the pair’s stage-1 score among the partners |  | • |  |

#### S5.2. Assembly Pseudocode

##### Algorithm S1

HARICOT

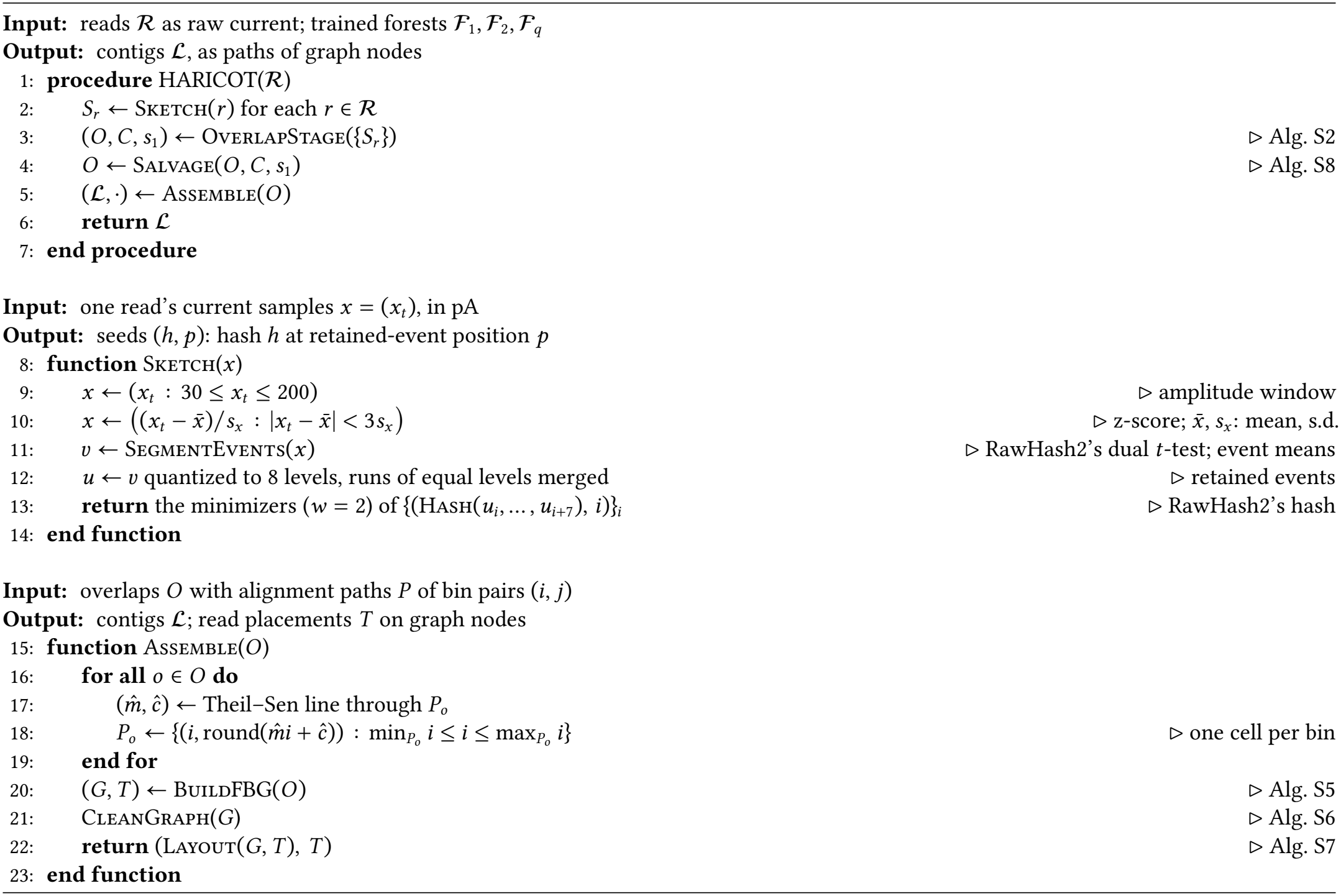

##### Algorithm S2

OverlapStage

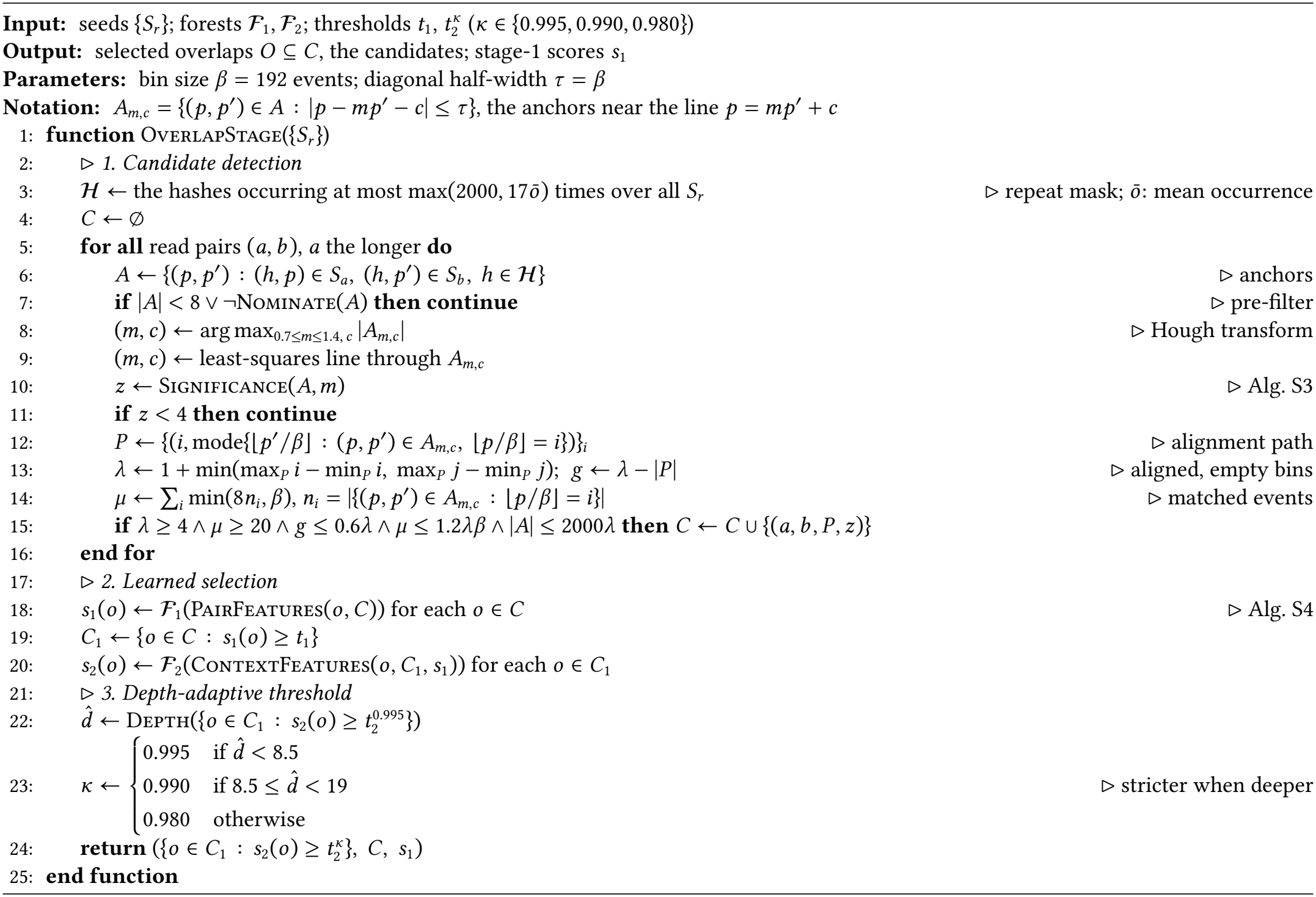

##### Algorithm S3

Significance, NullCount and Nominate

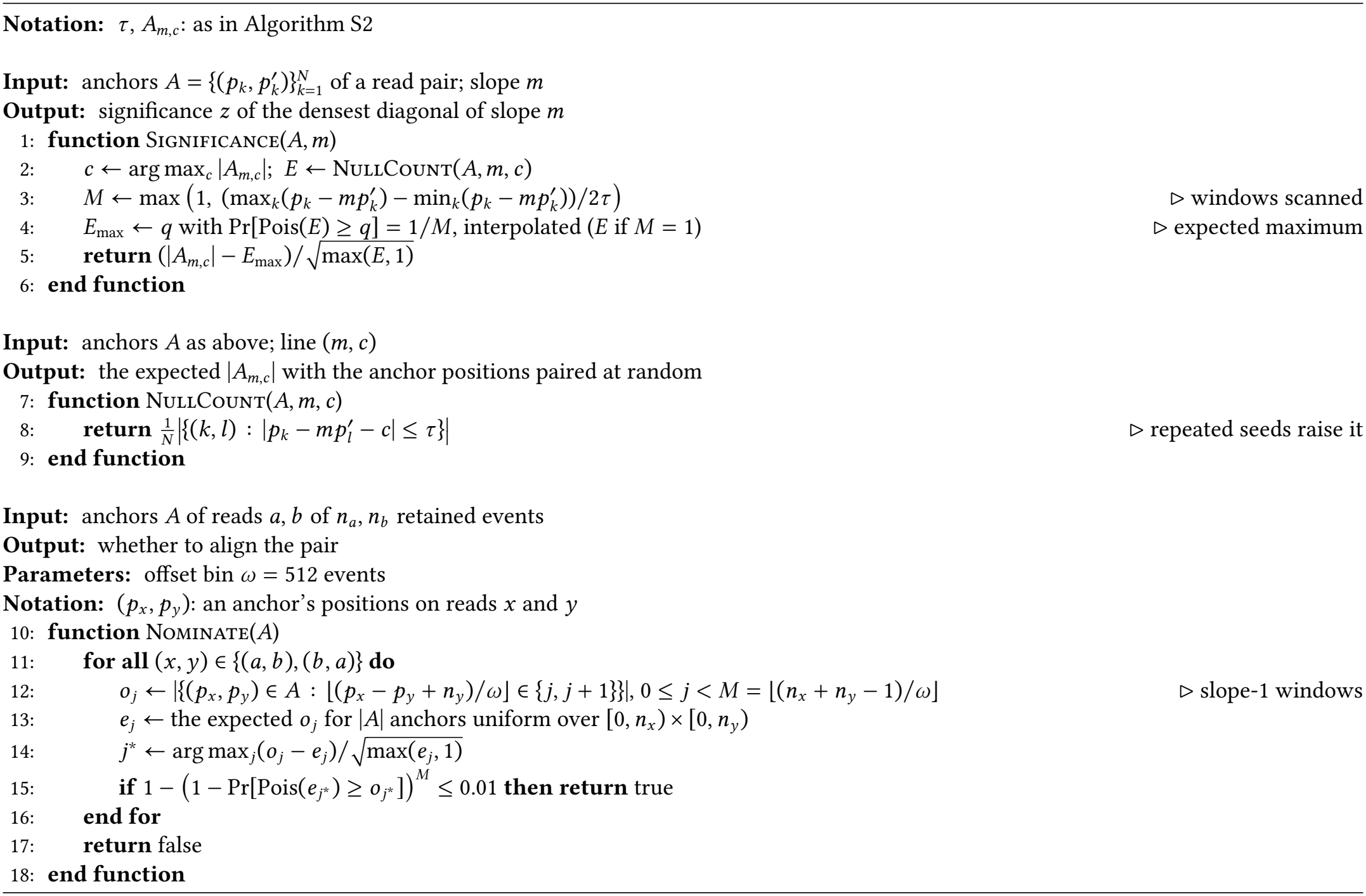

##### Algorithm S4

PairFeatures, ContextFeatures and Depth

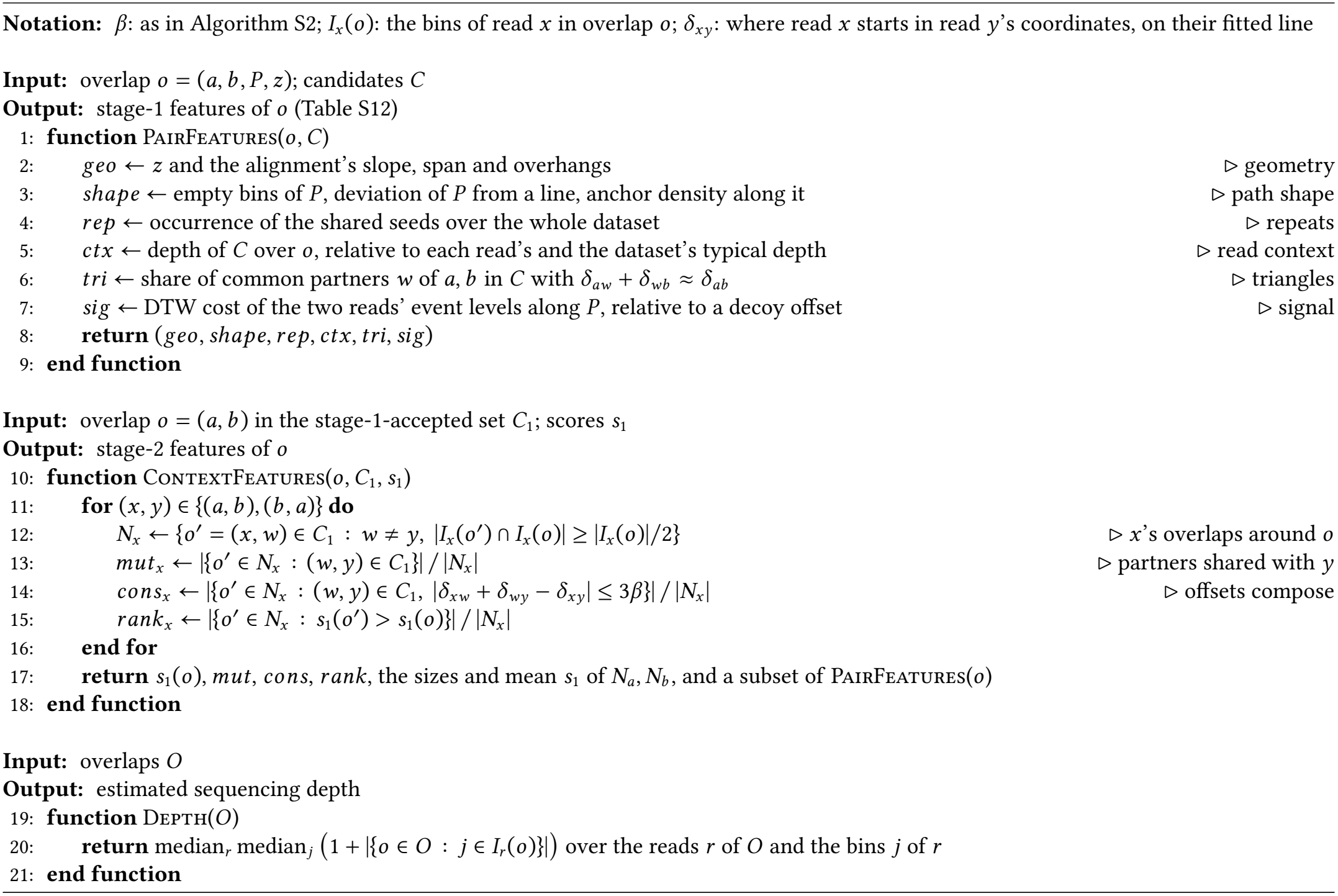

##### Algorithm S5

BuildFBG

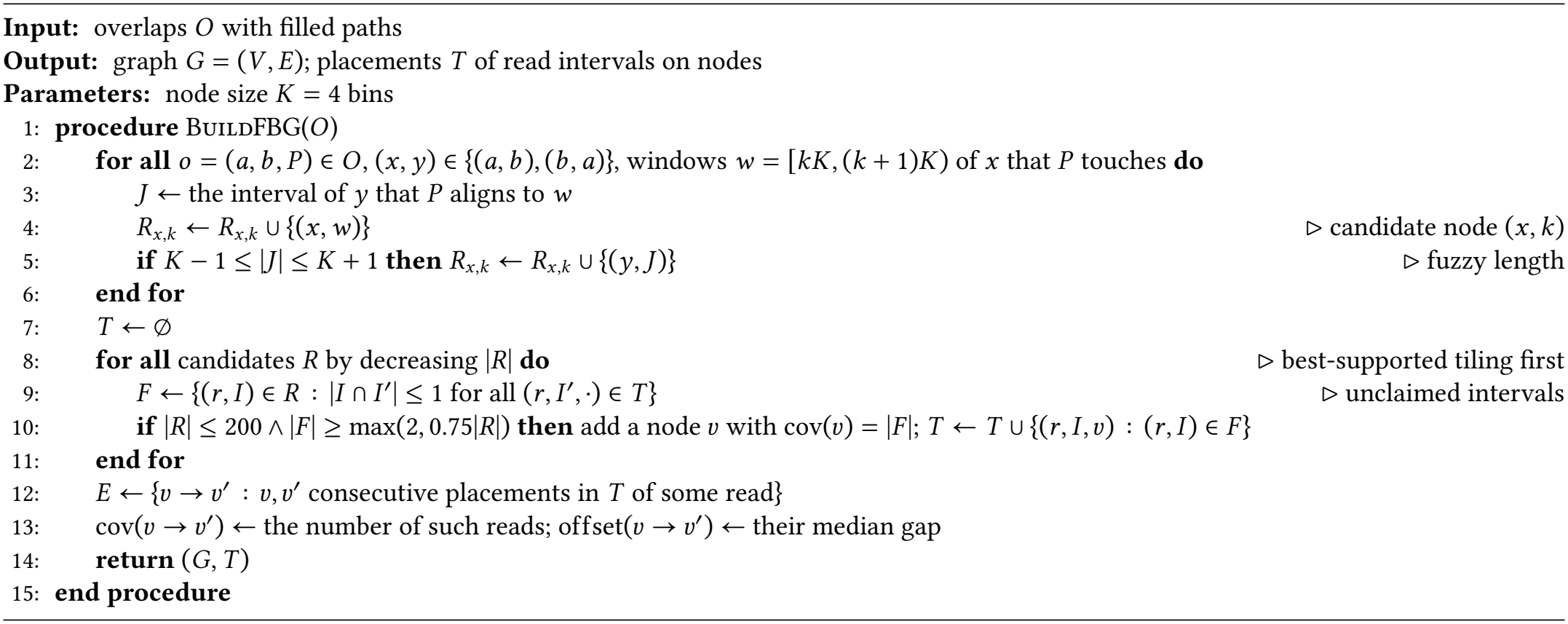

##### Algorithm S6

CleanGraph

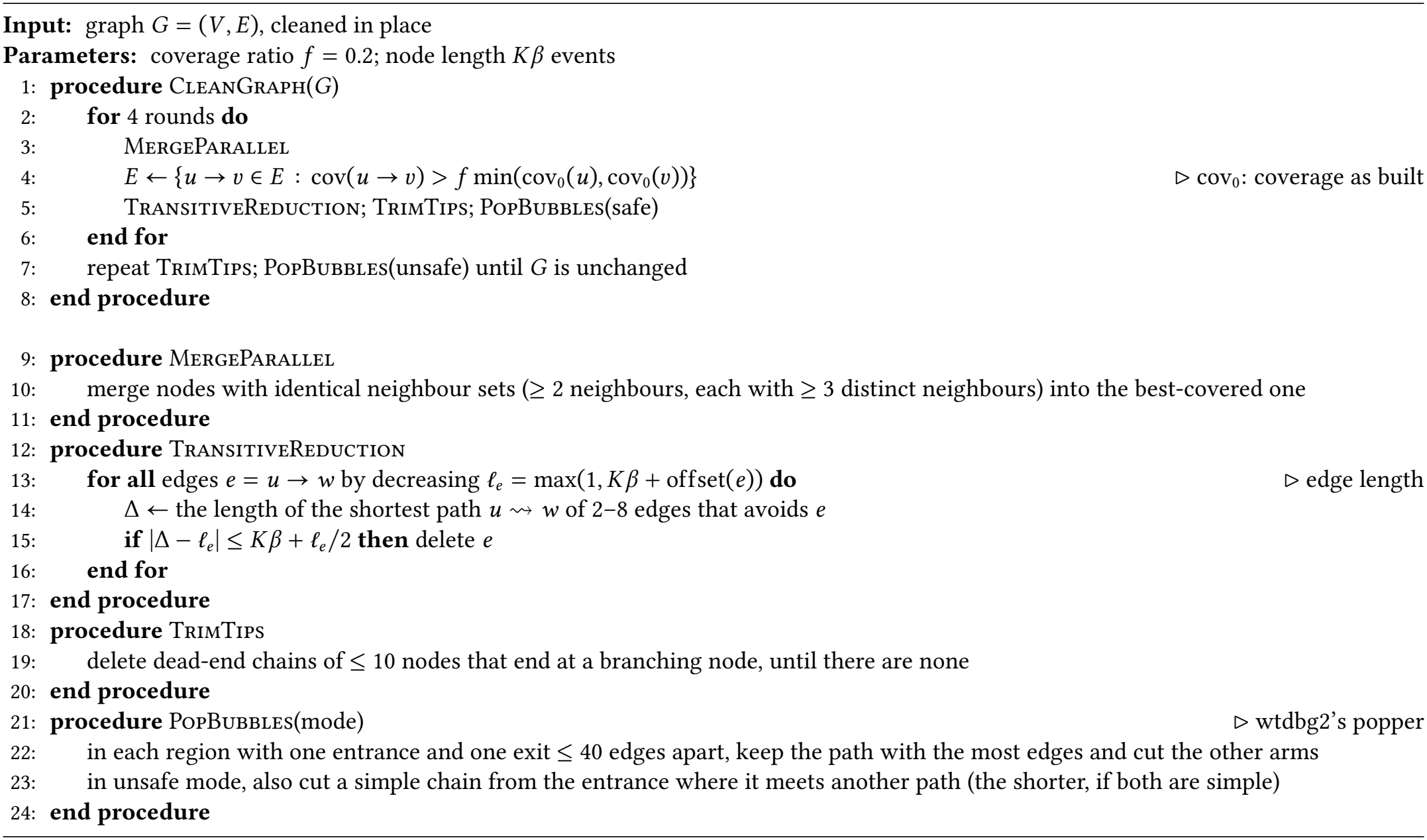

##### Algorithm S7

Layout

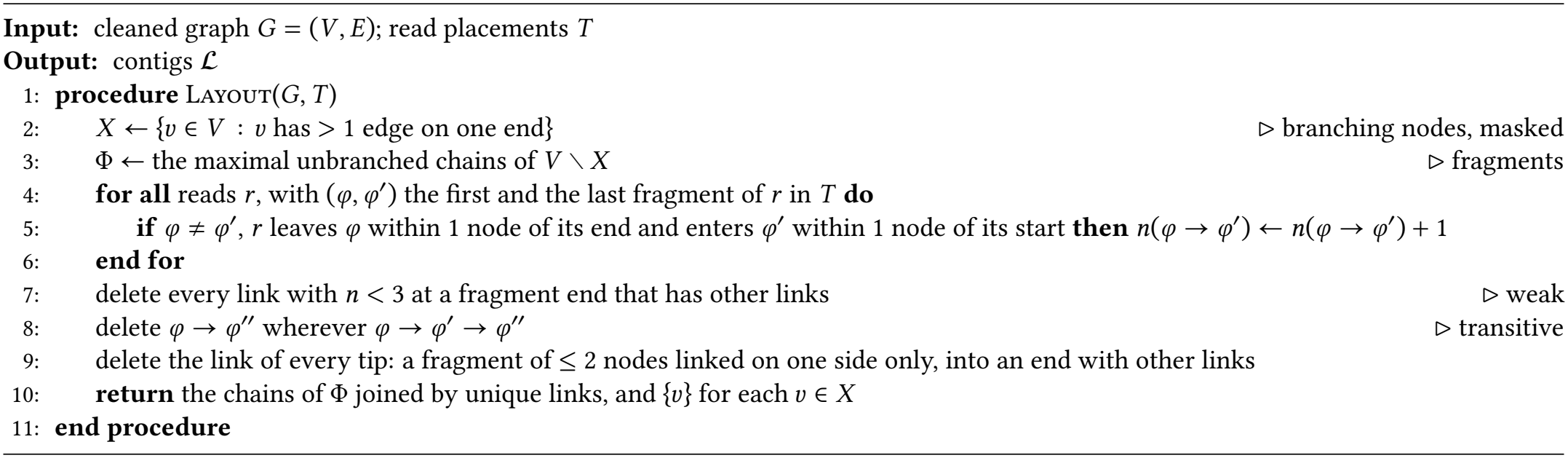

##### Algorithm S8

Salvage

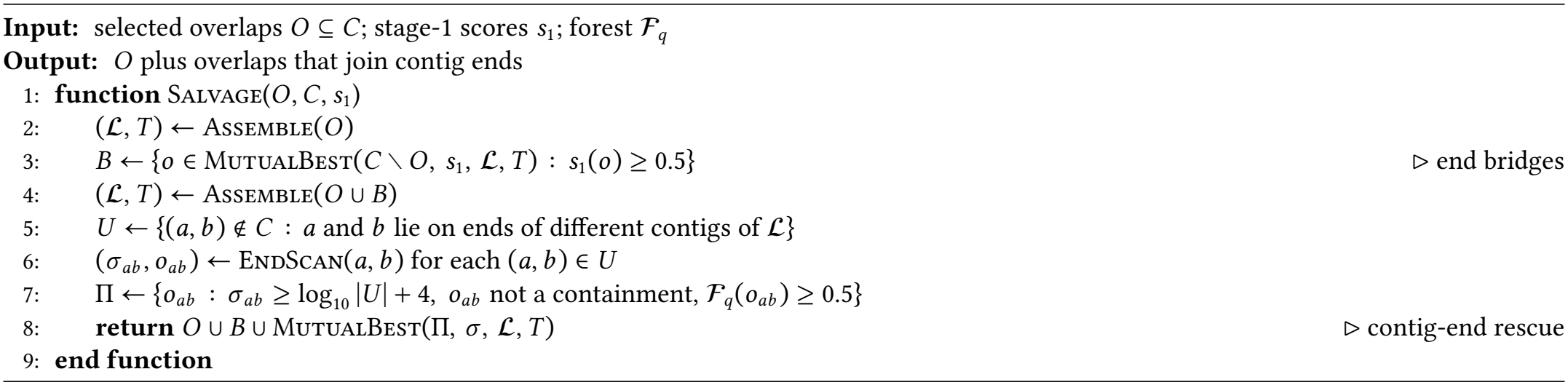

##### Algorithm S9

MutualBest and EndScan

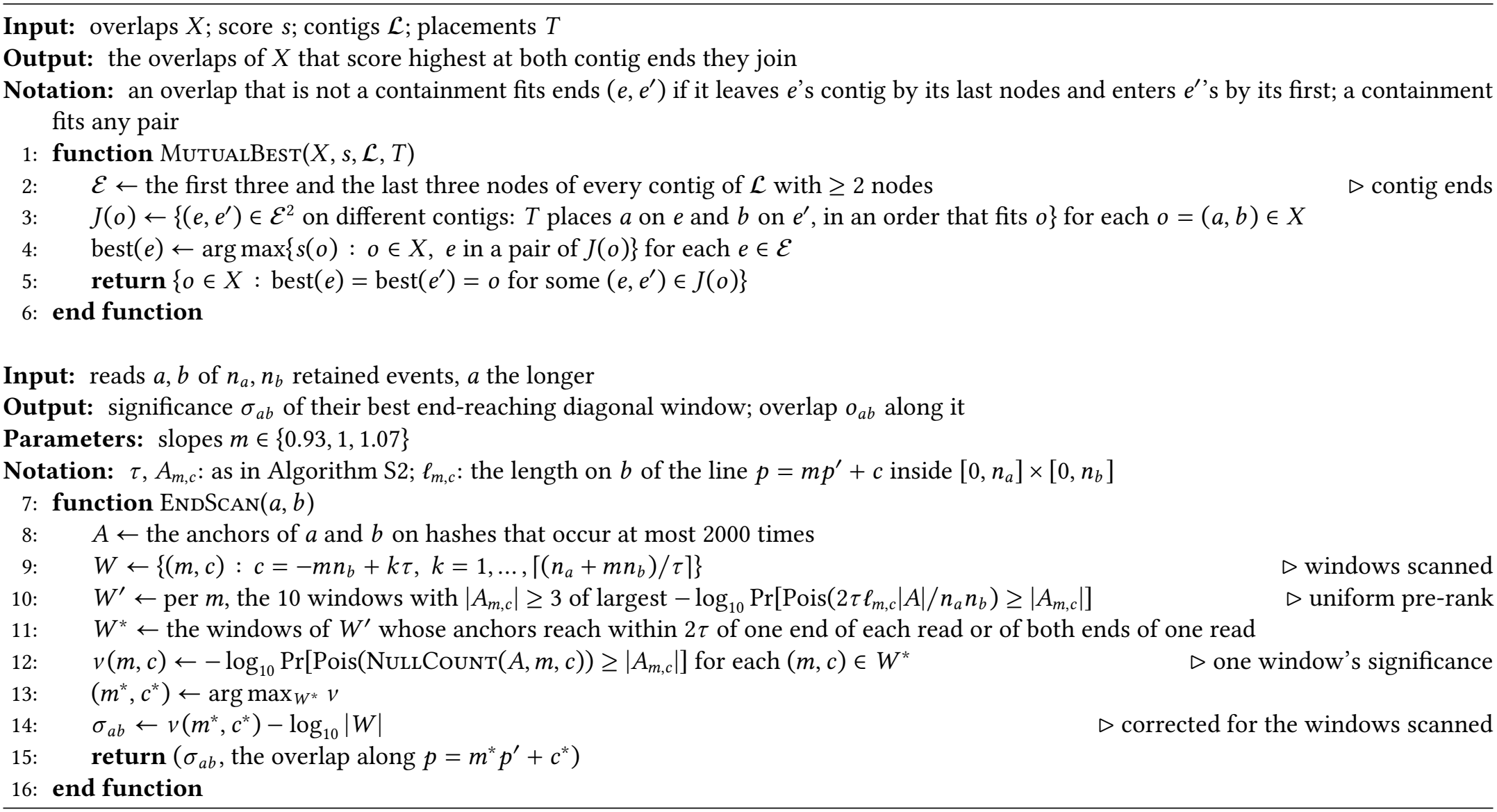

